# TI-Toolbox: An Open-Source Software for Temporal Interference Stimulation Research

**DOI:** 10.1101/2025.10.06.680781

**Authors:** Ido Haber, Aksel Jackson, Axel Thielscher, Aviad Hai, Giulio Tononi

## Abstract

**Background:** Temporal interference stimulation is a novel non-invasive brain stimulation approach that promises selective targeting of deep brain structures while minimizing off-target cortical stimulation. Despite a growing interest in temporal interference applications, there is a need for integrated computational tools that seamlessly connect neuroimaging data preprocessing through montage optimization, field simulation, and analysis within a unified framework designed for translational and clinical research.

**Methods:** We developed TI-Toolbox, an open-source software platform that integrates established neuroimaging tools (dcm2niix, SimNIBS, FreeSurfer) with specialized algorithms for temporal interference research. The platform provides end-to-end workflows encompassing structural MRI preprocessing, volume conduction modeling, montage optimization, electric field simulation, and region-of-interest analysis. Both graphical user interface and command-line interface implementations ensure accessibility across user expertise levels. The platform employs containerized deployment via Docker to ensure reproducibility and cross-platform compatibility.

**Results:** TI-Toolbox successfully automates the complete temporal interference research pipeline, from DICOM conversion through final field analysis. The platform demonstrates robust performance across operating systems and provides standardized workflows that enhance reproducibility. Furthermore, our case studies support the validity of our HD-EEG mapping approach for montage standardization and the need for individualized modeling for exposure assessment.

**Conclusions:** TI-Toolbox addresses critical infrastructure gaps in temporal interference research by providing researchers with a unified, validated platform that reduces technical barriers and accelerates translational research in non-invasive deep brain stimulation.

**Highlights:**

- Open-source platform unifying TI stimulation workflow end-to-end
- Docker deployment ensures reproducibility across operating systems
- HD-EEG electrode mapping preserves field characteristics
- Standardized model is sufficient for montage optimization
- Anatomical differences explain >40% of inter-individual variability

## 1. Introduction

Temporal interference (TI) stimulation has emerged as a promising technique for non-invasive neuromodulation, offering the potential to selectively target deep brain structures without surgical intervention (1). Unlike conventional transcranial electrical stimulation (tES) methods that predominantly affect superficial cortical regions, TI employs multiple pairs of high-frequency carriers with an offset frequency to create focal stimulation at depth through interference patterns (2). This approach addresses a fundamental limitation in non-invasive brain stimulation: the trade-off between stimulation depth and spatial focality.

The principle underlying TI stimulation involves the application of high frequency currents through multiple electrode pairs, typically using frequencies in the kilohertz band such as 2.0 kHz and 2.01 kHz, which generate an amplitude-modulated envelope at a physiologically relevant offset frequency (10 Hz in this example) (Fig 1) (3). The dominant working hypothesis in the field posits that while neurons cannot directly follow kilohertz oscillations due to their slower membrane time constants, they can respond to the interplay of the interfering fields through various proposed mechanisms including envelope demodulation, linear integration and signal mixing (4–6). Though there is not yet an established mechanistic explanation, recent preclinical studies have demonstrated successful modulation of hippocampal activity and motor cortex functions (7,8) while emerging human studies report effects on improvement of working memory, reduction of epileptic biomarkers, and enhancement of slow wave activity during sleep. (9–11).

**Figure 1.**
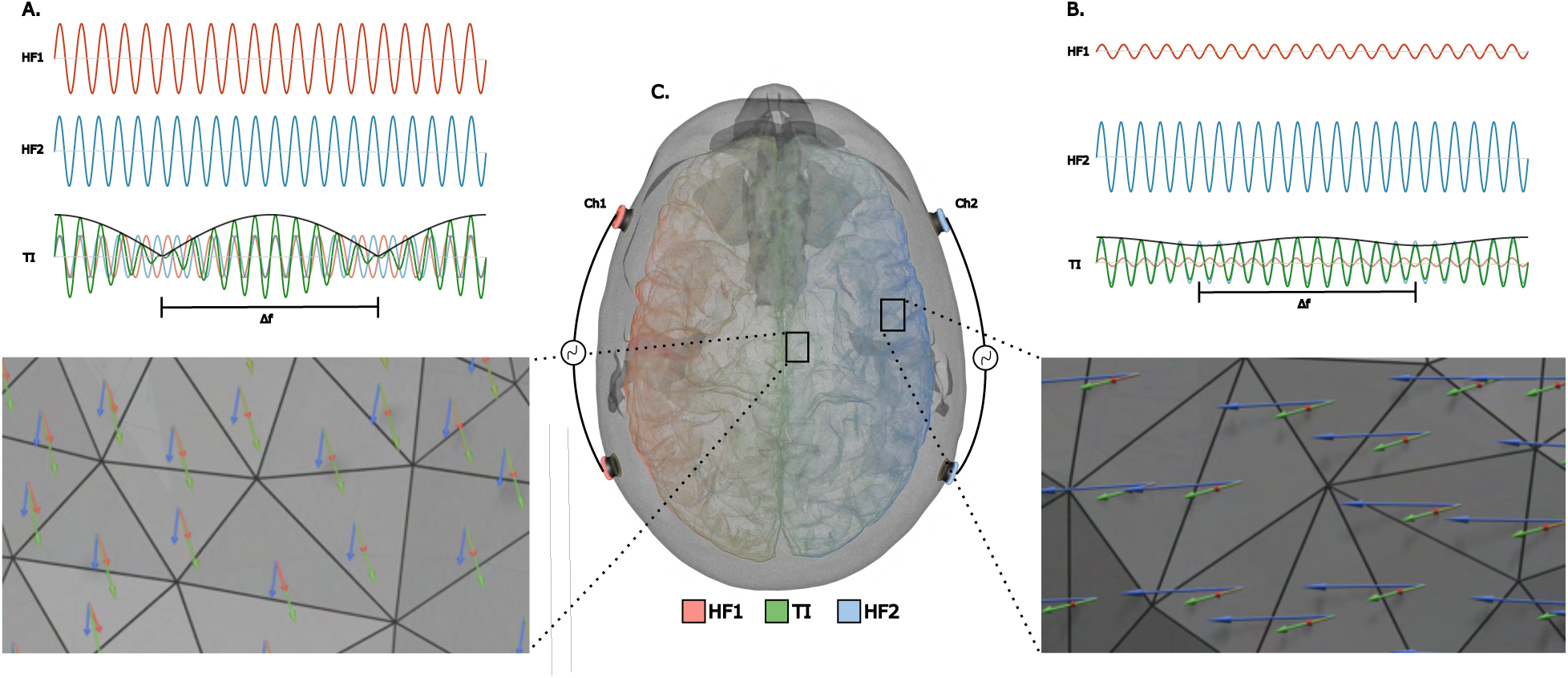
Temporal and spatial characteristics of temporal interference (TI) field distribution. **A.** Central brain region showing maximal modulation amplitude where TI envelope (Δf) dominates over high-frequency (HF) carrier components (HF₁,HF₂). **B.** Superficial cortical area demonstrating minimal modulation with dominant HF₂ carrier activity. **C.** Three-dimensional head model illustrating electrode montage with HF₁ (carrier frequency), HF₂ (carrier frequency + Δf), and resulting TI vectors displayed in distinct colors. Field vectors represent electric field direction and magnitude at surface of the gray matter.

While researchers are actively working to decipher the neural mechanisms of TI at the microscopic level, modulation of the central nervous system critically depends on the electric field strength and likely the direction experienced by neural tissue in line with more traditional TES interventions (12,13). Therefore, knowing the electric field exposure is essential for planning rigorous clinical studies. Some studies take advantage of depth electrodes for cohorts that have surgical implants, which allows field distribution to be assessed directly and tuned towards the desired target (10,14). However, in most human studies, participants do not have implanted electrodes, making it impossible to directly measure the brain’s exposure to TI stimulation.

To address this challenge, in-silico simulations offer a powerful, non-invasive solution which enables researchers to estimate field exposure across the entire head with high spatial resolution, under controlled and reproducible conditions (15). These simulations are well-suited for conducting experimental manipulations, optimizing stimulation protocols, and minimizing possible risk to participants (16,17). In practice, however, existing solutions remain fragmented, often requiring researchers to assemble custom pipelines, or proving inaccessible due to steep learning curves or high licensing costs (18). This fragmentation poses significant challenges for establishing standardized workflows and hinders the broader clinical application of in-silico TI modeling.

Here, we introduce TI-Toolbox, an open-source platform that offers a comprehensive, integrated, end-to-end solution designed to tackle the key challenges in clinical TI research. The platform combines established neuroimaging tools within a unified framework specifically designed for TI applications, offering automated workflows from raw MRI data through optimized stimulation protocols and detailed field analysis (Fig 2A-D). TI-Toolbox aims to promote access to advanced TI modeling by unifying preprocessing, optimization, simulation, analysis and visualization within a single framework that promotes computational ease of use, reproducibility, and methodological standardization across the research community.

**Figure 2.**
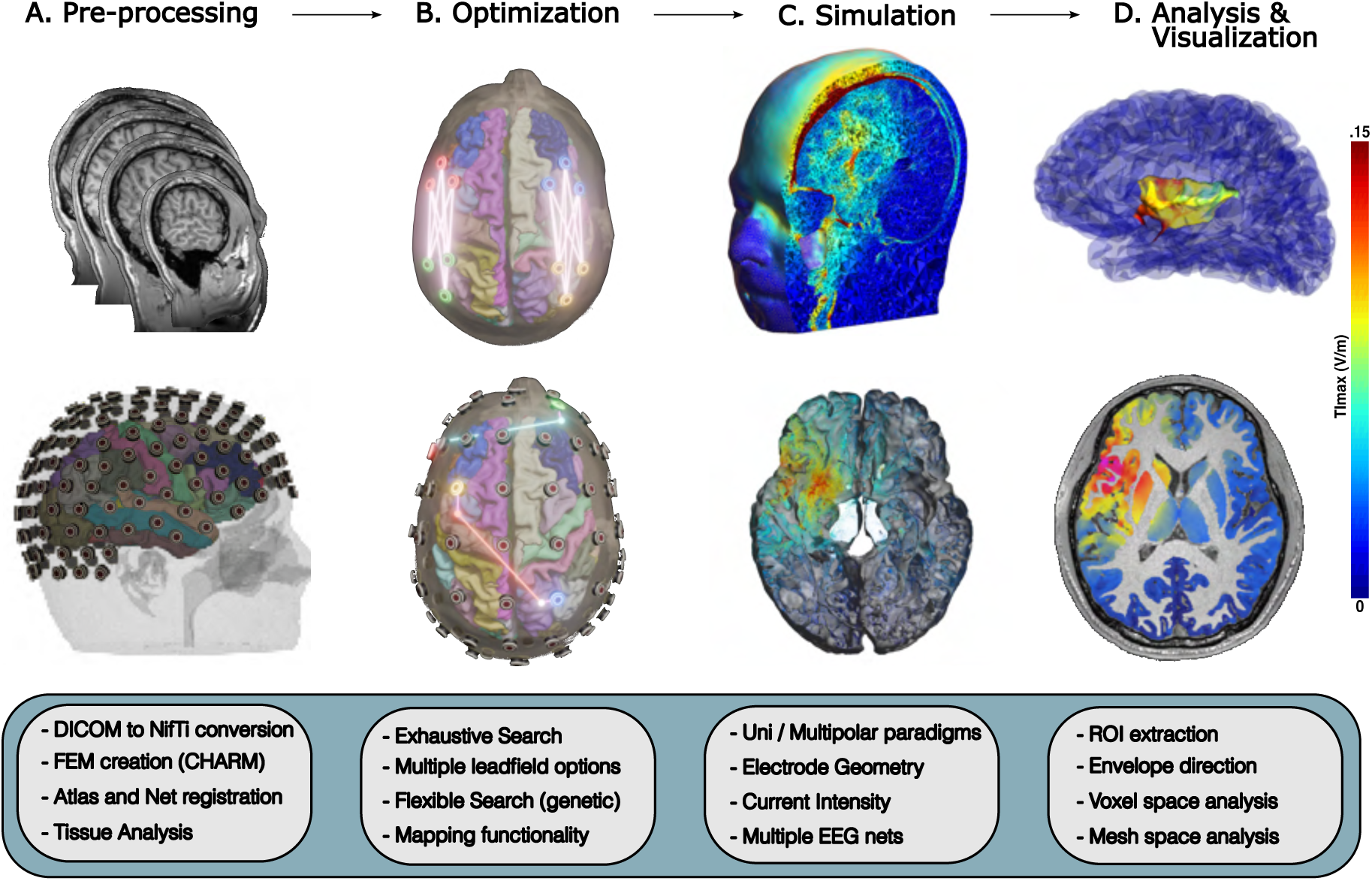
TI-Toolbox integrated computational pipeline. **A.** Preprocessing module: DICOM-to-NIfTI conversion via dcm2niix, FreeSurfer recon-all cortical reconstruction, and SimNIBS charm finite element method (FEM) head model generation. **B.** Optimization algorithms: flex-search genetic algorithm and exhaustive search approaches for electrode montage determination with region-of-interest (ROI) targeting. **C.** Simulation engine: magnitude and direction computation of TI_max_. **D.** Analysis and visualization: ROI extraction, and mesh/volumetric output generation compatible with Gmsh and Freeview.

Using TI-Toolbox we sought to answer multiple concerns that the field has been debating: (Q1) Does mapping optimized montages from an unconstrained genetic solution onto standard high-density EEG (HD-EEG) nets alter electric field characteristics (19,20)? (Q2) How does using individualized head models influence TI optimization performance versus a generalized model (21)? (Q3) Can demographic or anatomical factors explain inter-individual variability in TI exposure? To answer these questions, we evaluated three ROIs: the left insula, the right hippocampus, and a spherical ROI centered at MNI coordinates (36.10, 14.14, 0.33) with radius 5 mm, while assessing the intensity, direction and focality of the maximal modulation vector field.

## 2. Methods

### 2.1 Overview

The TI-Toolbox is organized into four main components: preprocessing, optimization, simulation, and analysis (Fig 2). The modules are designed to operate in a linear sequence, with standardized data structures and outputs enabling reproducibility across stages. The preprocessing module generates subject-specific head models from raw MRI data via automated DICOM conversion, surface reconstruction, and finite element meshing. The optimization module determines effective electrode configurations fitted to the subject anatomy and a stimulation objective, supporting both genetic and exhaustive algorithms. The simulation module computes subject-specific electric fields and TI envelopes, with support for directional field components, multi-polar TI, and customizable electrode configurations. The analysis module extracts region-wise metrics and generates visualizations through either surface (mesh) or volumetric (voxel) evaluation pipelines, with outputs viewable in Gmsh (22) (mesh) and Freeview (23) (voxel), and with optional cohort-level exports in MNI space.

Each component is accessible through both a command line interface (CLI) and a PyQt5- based graphical user interface (GUI). Containerization via Docker (24) ensures reproducibility (25) and simple orchestration of external dependencies across operating systems (Fig3A-C).

### 2.2 Prerequisites

To utilize TI-Toolbox, users must provide anatomical MRI data in DICOM format, with at least a T1-weighted sequence required for structural processing. The inclusion of T2- weighted images is recommended to improve tissue segmentation accuracy, and diffusion-weighted imaging (DWI) data may be supplied for the computation of directional conductivity tensors for running anisotropic simulations.

The platform is compatible with Windows, Linux, and macOS operating systems via Docker-based containers running Ubuntu. Docker (or Docker Desktop) is the only required local installation; all other dependencies are encapsulated within the containerized environment.

Access to the graphical user interface requires X11 forwarding across all operating systems. For standard usage, a minimum of 32 GB RAM is recommended. Installation instructions and a detailed usage guide are available in the project’s website (26).

To facilitate immediate use and learning, TI-Toolbox natively ships with the Ernie’s T1- weighted and T2-weighted scans from previous work (27) and MNI152 data (28).

Once prerequisites are installed, the correct BIDS formatting should be provided for the rest of the tools to work properly. Specifically, the user needs to set up their ‘sourcedatà sub-directory with either raw DICOMs or NIfTI files. The rest of the required files are generated automatically as the user moves through the toolbox (Fig 3 A-C).

**Figure 3.**
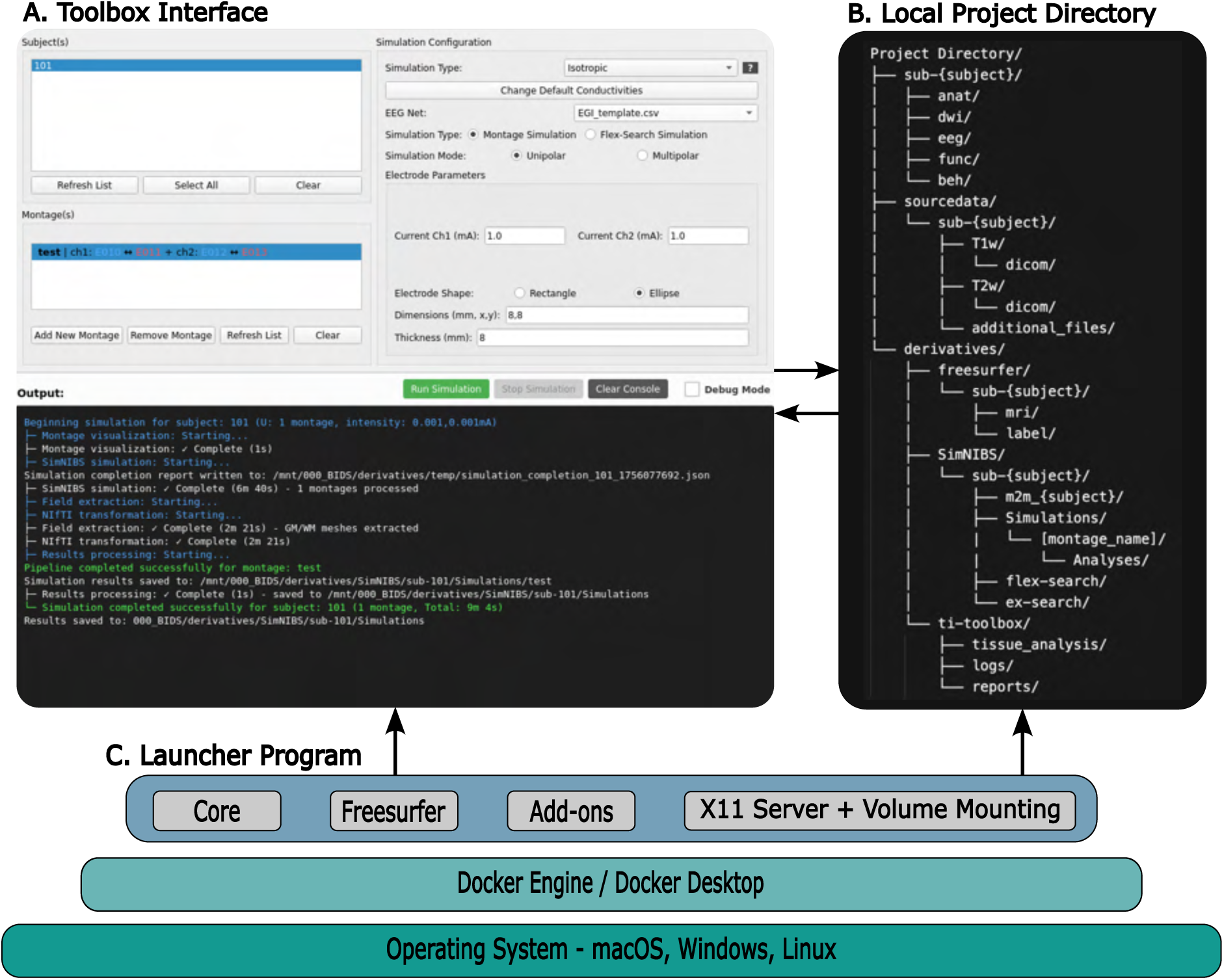
TI-Toolbox software architecture and containerized deployment. **A.** Unified interface providing graphical user interface (GUI) and command-line interface (CLI) access for workflow orchestration. **B.** Brain Imaging Data Structure (BIDS)-compliant project directory organization showing sourcedata/, derivatives/, and standardized file hierarchy. **C.** Multi-layered deployment architecture: host operating system supporting Docker containerization with launcher program managing SimNIBS and FreeSurfer dependencies through isolated container environments.

### 2.3 Core components and implementation

#### 2.3.1 Preprocessing pipeline

The preprocessing module orchestrates a workflow that transforms raw MRI data into simulation-ready head models through three primary stages: DICOM conversion, cortical reconstruction, and finite element method (FEM) creation.

DICOM to NIfTI conversion is performed using dcm2niix (29), which consolidates DICOM series into volumetric images. This process produces outputs compliant with the Brain Imaging Data Structure (BIDS) for compatibility with downstream workflows.

Structural processing leverages FreeSurfer’s ‘recon-all’ pipeline (23) through the ‘recon- all’ function, which performs cortical reconstruction including tissue segmentation, surface extraction, and automatic parcellation using standard atlases. The implementation supports both serial and parallel execution modes, with the parallel mode utilizing ‘GNU Parallel’(30) for efficient multi-subject processing.

Head model generation employs SimNIBS’s ‘charm’ function (31) for creating detailed FEMs. When both T1 and T2 images are available, charm utilizes multi-modal information for improved tissue segmentation accuracy. As part of ‘charm’ command, multiple 10-20 and high-density electroencephalogram (HD-EEG) nets are co-registered with the head model via non-linear transformation from Montreal Neurological Institute (MNI) to subject space (Fig 2A). All steps incorporate comprehensive error handling, generate timestamped logs, and sharable HTML reports (section 2.4).

Following FEM creation, a dedicated tissue analysis script (‘utils/tissue_analyzer.py’) estimates cortical bone morphology, ventricular and subarachnoid cerebrospinal fluid (CSF), and skin characteristics, producing subject-level metrics (Fig 7A,B). These metrics are exported alongside preprocessing reports and can serve as covariates for downstream inter-individual variability analyses (Section 2.6.5).

#### 2.3.2 Optimization algorithms

The optimization module implements two complementary approaches addressing the unique challenges of multi-electrode, multi-objective optimization in TI stimulation:

Flex-Search Algorithm (‘flex-search/flex-search.py’): Built upon SimNIBS’s ‘TesFlexOptimization’ class (32), this module implements an adaptive genetic algorithm. Flex-Search iteratively evolves electrode configurations by simulating montages, evaluating the resulting electric field in a specified ROI based on a user-defined goal, and mutating to improve solutions over generations. The core implementation centers around the ‘build_optimization()’ function, which configures optimization parameters including goal selection (mean field, maximum field, or focality), post-processing options (max_TI, dir_TI_normal, or dir_TI_tangential), and electrode configurations. To mitigate local optima, the tool supports a multi-start strategy in which the optimizer is launched multiple times with distinct seeds; users can specify the number of runs and automatically retain the best montage across starts. While the multi-start approach consistently improves targeting performance, the gains are typically modest (Figure S1).

The algorithm supports three distinct ROI definition methods through helper functions. The ‘_roi_spherical()’ function enables definition of spherical ROIs with customizable center coordinates and radii, ideal for targeting specific anatomical landmarks. The ‘_roi_atlas()’ function interfaces with cortical atlases (Desikan-Killiany, Destrieux, HCP-MMP1) (33–35) to target specific cortical regions by label. The ‘_roi_subcortical()’ function extends targeting capabilities to volumetric subcortical structures using ‘charm’ ‘Labeling.nii.gz’. The implementation also provides control over optimization hyper- parameters including maximum iterations, population size, and CPU core utilization for performance tuning.

The electrode mapping functionality (‘map_to_nearest_net_electrodes()’) utilizes an optimal assignment algorithm to project unconstrained optimization solutions onto standard EEG montages. The method constructs a Euclidean distance matrix between the optimized electrode positions and available positions from co-registered EEG nets. The Hungarian algorithm (via ‘scipy.optimize.linear_sum_assignment’) then solves the bipartite matching problem to minimize total assignment distance while ensuring a one-to-one mapping between optimized and standard positions (Fig 4A) (Table S1) (Fig. S5).

**Figure 4.**
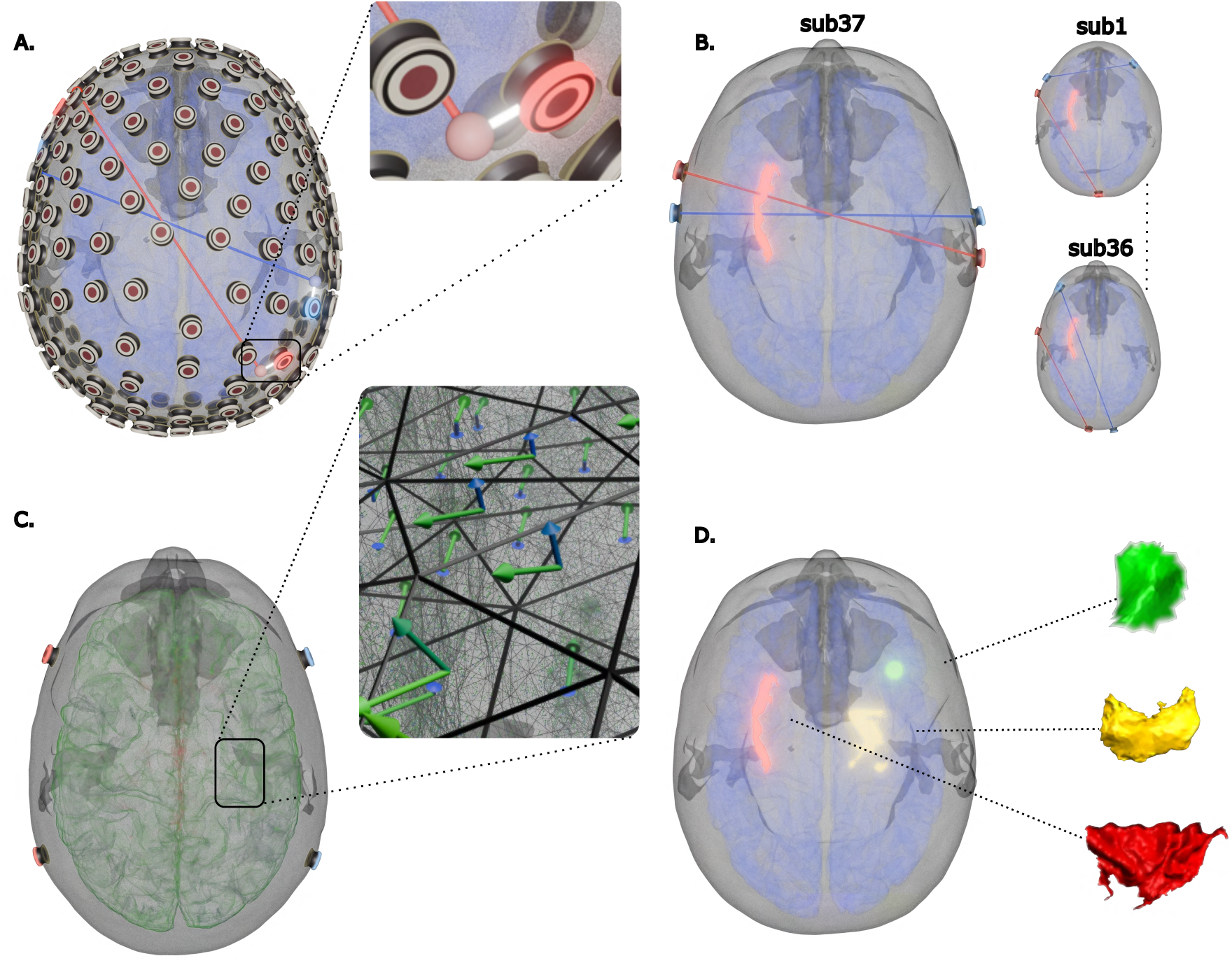
Target definition and optimization objectives for case studies. **A.** Electrode mapping functionality demonstrating unconstrained optimization to standard high-density electroencephalography (HD-EEG) net projection using Hungarian algorithm for optimal electrode assignment. **B.** Head model comparison between subject-specific anatomical models (subjects 1-36) and generalized template (Ernie model, subject 37). **C.** Field component illustrations: TI_max_ (maximal modulation) and TI_normal_ (surface-aligned normal component) vectors relative to cortical surface geometry. **D.** Target regions: left insula (cortical, red), right hippocampus (subcortical, yellow), and spherical ROI (green) at Montreal Neurological Institute (MNI) coordinates (36.10, 14.14, 0.33) with 5 mm radius, representing diverse anatomical targeting scenarios.

Exhaustive Search Algorithm (‘ex-search/ti_sim.py’): This module relies on a pre- computed leadfield and performs systematic evaluation of configurations through cartesian product operations, ensuring a logical balance between coverage of the search space and compute efficiency (Fig 2B top).The ex-search leverages SimNIBS’s ‘TI_utils.py’ module for efficient field calculations. The ‘load_leadfield()’ function retrieves pre- computed electromagnetic field distributions, while ‘get_field()’ calculates the electric field for specific electrode configurations. Results are stored as mesh files with associated visualization configurations, enabling immediate inspection in Gmsh as well as a .csv file that includes intensity and focality metrics for the user to inspect (Table S2).

#### 2.3.3 Simulation engine

The simulation module (‘simulator/main-TI.sh’) offers a robust control over simulation parameters which interfaces with SimNIBS’s ‘sim_struct.SESSION’ class to configure parameters like anisotropy type (scalar, volume normalized, directional), electrode geometries, and current delivery.

For TI calculation, the module leverages SimNIBS internal functions to compute individual electric fields for each carrier frequency pair. These fields are then processed through the ‘TI_utils.get_maxTI()’ function, which implements the framework suggested by Grossman et. al. (1).

In addition to TI_max_, a surface-aligned normal component (TI_normal_) is computed from the local TI vector onto the middle layer of the gray-matter surface (Fig 4C). Both TI_max_ and TI_normal_ are exported and analyzed downstream as part of the analyzer tool.

Output management includes generation of both volumetric and surface-mapped results, with automatic conversion to MNI space for group-level analyses and visualizations. The module creates visualization files compatible with Gmsh and Freeview, facilitating immediate inspection of simulation results in mesh and voxel spaces.

Additionally, the platform implements multi-channel temporal interference approach (‘simulator/main-mTI.sh’) for configurations that involve more than two electrode pairs (multi-polar TI) (36) This extension supports emerging TI paradigms that utilize more than two channels to achieve enhanced spatial selectivity, steering, or multi-target stimulation (Figure S2).

For reproducibility and batch processing, all electrode montages are saved to and loaded from a centralized montage database (‘montage_list.json’) that maintains both unipolar and multipolar montage definitions for various EEG cap systems.

High-Frequency Field Assessment: The simulator outputs detailed analysis of the individual carrier fields before interference calculation. A summary statistics file (‘fields_summary.txt’) documents key metrics including mean, maximum, and percentile values for each carrier field within gray matter. Specific Absorption Rate (SAR) can be derived from this output according to the safety recommendations from Cassarà et al. 2025 (16,17) which will be integrated in future version of the TI-Toolbox.

All outputs maintain consistent naming conventions incorporating subject identifiers, target names, and field types, facilitating automated processing in downstream pipelines. The modular output structure supports selective export based on computational resources and research needs, with options to disable specific output types through configuration parameters.

#### 2.3.4 Analysis and visualization

The analysis module (‘analyzer/mesh_analyzer.py’) provides tools for quantifying and visualizing TI stimulation outcomes through the ‘MeshAnalyzer’ class. This class implements three primary analysis modes: spherical ROI analysis, cortical region analysis, and whole-head analysis.

The ‘analyze_sphere()’ method enables analysis of spherical regions of interest, utilizing SimNIBS’s mesh manipulation functions to extract field values within specified coordinates and radii. For cortical analysis, the ‘analyze_cortex()’ method interfaces with multiple anatomical atlases through SimNIBS’s ‘subject_atlas()’ function, supporting Desikan-Killiany, Destrieux, and HCP-MMP1.

A surface mesh generation pipeline implemented through the ‘_generate_surface2_mesh()’ method, which calls the ‘msh2cortex’ utility to project volumetric field data onto the middle layer of the cortex. This enables accurate analysis of field distributions in gray matter while accounting for cortical folding patterns (Fig 2D top).

The ‘analyze_whole_head()’ method performs evaluation across all regions in a specified atlas, generating both region-specific analyses and population-level summaries. Field statistics are computed using weighted averaging based on node areas obtained through the ‘nodes_areas()’ function, ensuring accurate representation of field distributions on irregular meshes.

Visualization capabilities are implemented through the ‘MeshVisualizer’ class (‘analyzer/visualizer.py’), which generates multiple output formats including 3D mesh visualizations with customized colormaps (viewable in Gmsh), ROI weighted field distribution histograms, and region-wise scatter plots. The ‘_generate_region_visualization()’ method creates masked field displays highlighting specific anatomical regions, while ‘_generate_whole_head_plots()’ produces summary visualizations for multi-region analyses.

For volumetric analysis, the ‘VoxelAnalyzer’ class (‘analyzer/voxel_analyzer.py’) provides complementary functionality for analyzing field distributions in NIfTI format. This includes the ‘extract_values_from_roi()’ method for spherical and atlas-based ROI extraction which can be inspected in Freeview (Fig 2D bottom).

Group-level analysis is implemented via the ‘group_analyzer.py’, which enables collection and comparison of TI simulation results across multiple subjects. This module generates averaged volumetric field distributions in NIfTI format, producing inter-subject comparison plots, and exporting summary statistics across the cohort. Group level analysis remains in subject space for cortical and sub-cortical regions while arbitrary spherical targets are defined in MNI space and automatically transform into each subject’s native space using SimNIBS ‘mni2subject_coords’ method.

### 2.4 Standardization and reproducibility

TI-Toolbox aims to ensure complete reproducibility and standardization across clinical research workflows. The platform’s logging infrastructure, built on the centralized ‘logging_util’ module, provides consistent formatting and hierarchical logging across all components. The ‘get_logger()’ function creates timestamped log files following the pattern ‘[YYYY-MM-DD HH:MM:SS] [module_name] [level] messagè, ensuring precise tracking of all operations.

The ‘configure_external_loggers()’ function extends this logging consistency to external dependencies including SimNIBS, and FreeSurfer, redirecting their outputs through the platform’s unified logging system. This integration ensures that all processing steps, regardless of their origin, are captured in a single, searchable log file stored in the BIDS-compliant (37) ‘ derivatives/ti-toolbox/logs/sub-{SUBJECT_ID}/’ directory structure.

HTML report generation is managed through the ‘report_util’ module, which provides two specialized generators: ‘PreprocessingReportGenerator’ and ‘SimulationReportGenerator’. These generators create comprehensive reports that summarize key processes and outcomes like fMRIPrep (38), documenting aspects of the pipeline. The ‘create_preprocessing_report()’ function captures structural processing steps, quality metrics, and warning/error logs, while ‘create_simulation_report()’ documents electrode configurations, stimulation parameters, and field calculation results.

### 2.5 Deployment and Infrastructure

#### 2.5.1 Docker Compose

TI-Toolbox’s deployment architecture leverages Docker Compose orchestration to manage a complex ecosystem of neuroimaging tools while ensuring consistent behavior across diverse computing environments. The ‘docker-compose.yml’ configures the primary service containers: Core (+SimNIBS), FreeSurfer, connected through a dedicated bridge network (‘ti_network’) (Fig 3C).

Volume management is handled through named Docker volumes that persist software installations across container restarts, significantly reducing startup times. The ‘LOCAL_PROJECT_DIR’ environment variable enables flexible data mounting, allowing users to process data stored anywhere on their filesystem while maintaining isolation between the host and container environments.

#### 2.5.2 Launcher program (Bash / Executable)

The toolbox implements several deployment strategies to accommodate different use cases:

Desktop Deployment: The executable launcher (‘launcher/executable/’) provides platform-specific binaries created through PyInstaller (39). The executables implement a PyQt5-based GUI (40,41) that manages Docker daemon connectivity, container lifecycle, and volume mounting. The launcher includes error handling through the ‘dialogs.py’ module, providing user-friendly feedback for common issues such as Docker daemon availability, disk space, and permission errors. The executable requires no system installation after downloading from either the GitHub release page or the website’s release page (22).

Server Deployment: The bash-based launcher (‘launcher/bash/loader.sh’) implements a deployment script that handles the setup necessary for healthy operation of the toolbox. The bash entry point provides a solution for headless servers processing and can be used as a desktop alternative to the executable approach.

#### 2.5.3. Batch processing and parallelization

TI-Toolbox implements batch processing and parallelization strategies to enable efficient analysis of large clinical cohorts. The platform’s parallelization architecture operates at multiple levels, from individual processing steps to cohort-wide analyses.

Subject-Level Parallelization: The preprocessing pipeline (‘pre-process/structural.sh’) implements GNU Parallel integration (25) for concurrent processing of multiple subjects. When invoked with the ‘--parallel’ flag, the system automatically detects available CPU cores and distributes subjects across parallel workers. The implementation includes intelligent load balancing through the ‘parallel’ command’s built-in job scheduling.

Algorithm-Level Parallelization: The flex-search optimization module supports multi-core execution through the ‘--cpus’ parameter, which is passed directly to SimNIBS’s differential evolution optimizer for faster processing.

Pipeline-Level Batch Processing: The GUI employs multi-threading to keep the interface responsive while long-running jobs are carried out in background worker threads. A job queue coordinates preprocessing, optimization, simulation, and analysis tasks, with real-time progress reporting, log streaming, and graceful cancellation. This design allows concurrent execution of independent stages across subjects while preserving deterministic logging and outputs.

TMUX Parallelization: The code container which is exposed to the user interaction comes with TMUX (42) installed which enables further multiplexing where the user can span multiple instances of the toolbox simultaneously.

System Monitoring: To monitor the system during high-demand processes, a designated ‘System Monitor’ tab is available that tracks CPU and memory usage of the toolbox processes in real time.

### 2.6 Case Studies

#### 2.6.1 Targets and goals

Using the flex-search optimization algorithm, we evaluated three distinct brain targets (Fig 4D) with specific optimization objectives:

Left insula (deep cortical target): We performed two optimization rounds: (1) TI_max_ optimization maximizing the mean field intensity within the ROI, and (2) TI_normal_ optimization maximizing the mean surface-aligned normal component within the ROI. Both approaches focused on mean rather than maximum values to promote homogeneous field distributions.

Right hippocampus (subcortical target): We optimized for TI_max_, maximizing the mean field intensity within the ROI. Given that it is a sub-cortical target, we focused solely on intensity optimization without additional directional constraints.

Spherical ROI (MNI coordinates 36.10, 14.14, 0.33; radius 5 mm): We implemented three distinct optimization strategies: (1) mean optimization maximizing the mean TI_max_ intensity within the ROI, (2) focality optimization maximizing the ratio between mean TI_max_ in the ROI versus mean TI_max_ in gray matter using various thresholding approaches, and (3) max optimization maximizing the maximum value of the TI_normal_ component within the ROI.

#### 2.6.2 subjects

We collected magnetic resonance imaging (MRI) scans of thirty-six participants (mean age = 29.9 ± 9.7 years, 58.3% Female) from the STRENGTHEN clinical trial (Table S4). Participants were excluded if they had any neuroradiologist-identified brain structural abnormalities. All participants gave written informed consent in accordance with the University of Wisconsin-Madison Institutional Review Board. MRI data were collected using a 3 Tesla MAGNUS (Microstructure Anatomy Gradient for Neuroimaging with Ultrafast Scanning, GE Healthcare) head-only MRI scanner. Structural images were acquired using T1- and T2-weighted images, with 0.8 mm isotropic voxels, and the following parameters.

T1-weighted: sequence = MP-RAGE (Magnetization-Prepared Rapid Gradient-Echo), repetition time (TR) = 2000 ms, echo time (TE) = 3 ms, inverse time (TI) = 1100 ms, flip angle = 8 degrees, field of view (FOV) = 256 x 256 mm2, matrix size = 320 x 320 pixels, resolution = 0.8 mm x 0.8 mm x 0.8 mm, number of slices = 240, acquisition time = 4 min. T2-weighted: sequence = CUBE-T2, TR = 2500 ms, TE = 90 ms, echo train length (ETL) = 120, FOV = 256 x 256 mm2, matrix size = 320 x 320 pixels, resolution = 0.8 mm x 0.8 mm x 0.8 mm, number of slices = 240, acquisition time = 4 min. Our 37th participant which serves as our ‘generalized model’ is the publicly available Ernie model (23).

#### 2.6.3 Flow

Raw DICOM files were preprocessed as in Section 2.3.1. For each target and objective, we ran the flex-search optimizer with the ROI definitions in Section 2.6.1, using a three-run multi-start configuration and retaining the best solution per subject (configurable via Advanced Settings: Number of Optimization Runs). For each participant, we then simulated the resulting montages with default isotropic conductivities (43–45) (Table S3), 1 mA per channel, 8 mm electrode diameter, 4mm thick saline gel and a 2mm thick rubber electrode on top. Finally, we applied group analysis to extract TI_max_, TI_normal_, and focality within ROIs and gray matter.

#### 2.6.4 Statistical analysis

Statistical analyses were conducted using Python with scipy, statsmodels, and pandas libraries. All analyses were performed using a two-tailed significance level of α = 0.05. Data normality was assessed using the Shapiro-Wilk test (‘scipy.stats.shapiro’), with parametric (paired t-test via ‘scipy.stats.ttest_rel’) or non-parametric (Wilcoxon signed-rank test via ‘scipy.stats.wilcoxon’) approaches selected based on normality test results (p > 0.05 indicating normal distribution).

For comparison analyses (Q1 and Q2), paired statistical tests were used to evaluate differences between conditions within subjects. Effect sizes were calculated using Cohen’s d for parametric tests (computed as mean difference divided by pooled standard deviation) and r (Z/√N) for non-parametric tests, where Z-scores were obtained from the Wilcoxon test statistic. Percentage changes were computed as ((B - A) / A) × 100, where A and B represent the two conditions being compared.

For inter-individual variability analyses (Q3), we investigated anatomical determinants of TI field exposure using data from the optimized electrode montages. First, we assessed multicollinearity among anatomical predictors (age, bone thickness/volume, CSF thickness/volume, and skin thickness/volume) using correlation analysis. Based on high correlations within tissue types, we selected volume measures over thickness to reduce avoid multicollinearity while retaining predictive information.

We then employed multiple linear regression (MLR) models constructed using ‘statsmodels.api.OLS’ with ordinary least squares fitting to quantify the overall variance explained by anatomical factors (age, CSF volume, bone volume, and skin volume) in predicting TI field intensity. Model fit was evaluated using R², adjusted R², and F-statistics. The models were applied separately to the left insula and right hippocampus targets.

## 3. Results

### 3.1 Overview

We investigated TI field characteristics across three distinct brain targets in a cohort of 36 participants, using our flex-search optimization approach. For the left insula (deep cortical target), we evaluated solutions from both TI_max_ optimization (maximizing mean field intensity) and TI_normal_ optimization (maximizing mean surface-aligned normal component). The right hippocampus (subcortical target) was analyzed using TI_max_ optimization focused on maximizing mean field intensity. The spherical ROI (MNI coordinates 36.10, 14.14, 0.33; radius 5 mm) was evaluated using three optimization strategies: mean TI_max_ intensity, focality (ratio of mean TI_max_ in ROI to gray matter), and maximum TI_normal_ component. All statistical comparisons employed paired tests selected based on normality assessments (two-tailed α = 0.05), with effect sizes and percentage changes calculated to evaluate both statistical and practical significance.

### 3.2 Mapping optimized solutions to standard HD-EEG montage

To assess whether constraining electrode placements to standard HD-EEG positions compromises optimization benefits, we compared field characteristics between freely optimized montages (optimized) and their mapped counterparts (mapped) across all three targets (n = 37, including the generalized model subject). These analyses focus on the mean optimization results—maximizing mean TI_max_ intensity for the left insula, right hippocampus, and spherical ROI.

For the left insula, mapping to standard electrode positions resulted in minimal reduction in field intensity, with mean TI_max_ decreasing from 0.324 V/m to 0.319 V/m (Δ = -0.0043 V/m, -1.35%, p = 0.010). The maximum field values showed similar small reductions from 0.470 V/m to 0.464 V/m (Δ = -0.006 V/m, -1.28%, p = 0.046). Notably, focality remained essentially unchanged (1.410 vs 1.412, p = 0.609), suggesting that spatial selectivity was preserved (Fig 5 A).

**Figure 5.**
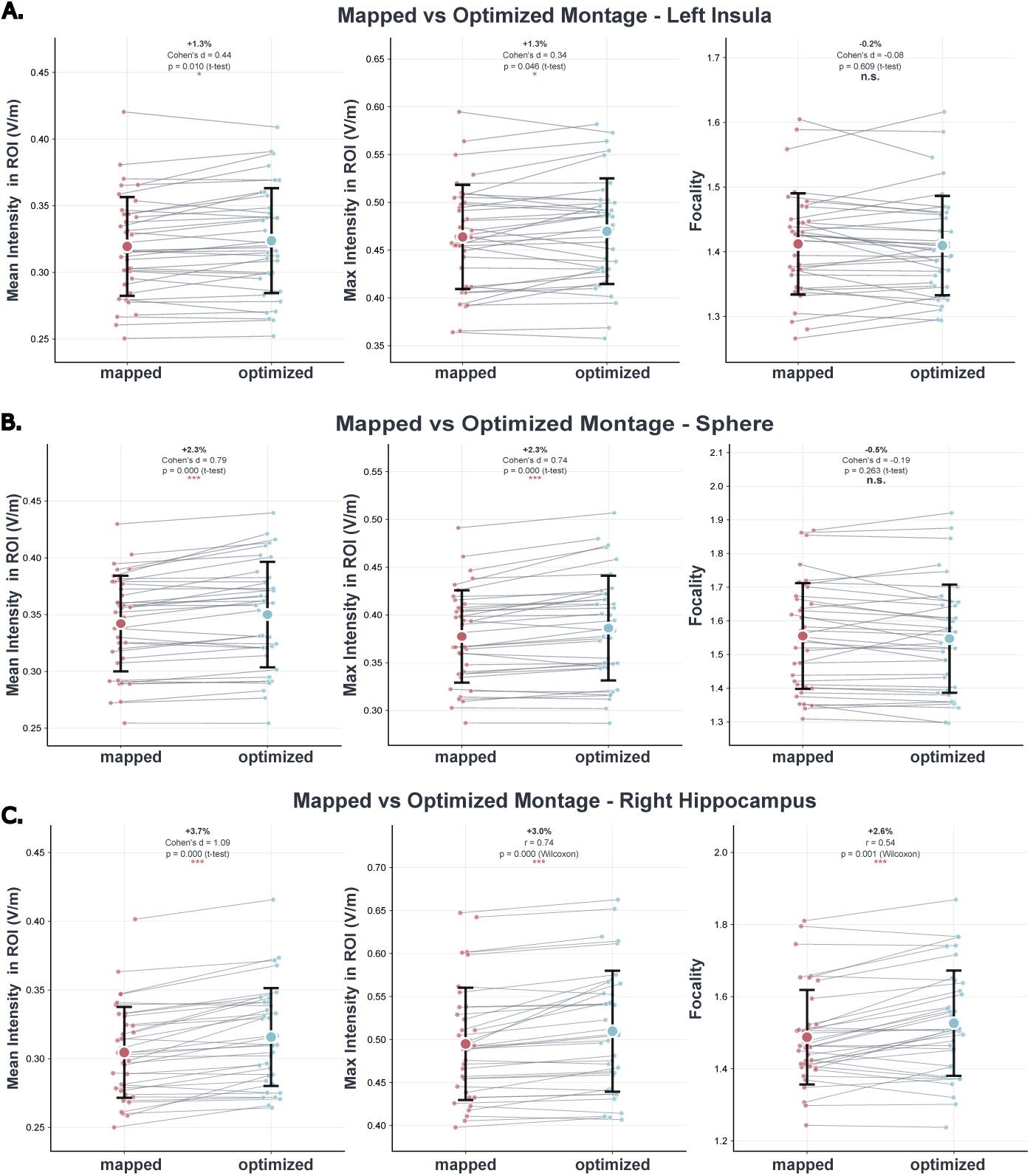
Comparison of optimized versus mapped electrode configurations. **A.** Within-subject comparison for left insula of TI_max_ characteristics between ‘mapped’ and ‘optimized’ conditions: mean TI_max_ = 0.324 vs 0.319 V/m (Δ = -1.35%, p = 0.010), maximum TI_max_ = 0.470 vs 0.464 V/m (Δ = -1.28%, p = 0.046), focality preserved (p = 0.609). **B.** Spherical target results: mean TI_max_ reduction of -2.31% (p = 2.66×10⁻⁴), maximum TI_max_ reduction of -2.33% (p = 6.66×10⁻⁴), focality unchanged (p = 0.263). **C.** Right hippocampus analysis: mean TI_max_ reduction of -3.67% (p = 1.07×10⁻⁷), focality reduction of -2.62% (p = 0.000642). Individual participants are shown as small colored dots (n=37) with slight horizontal jitter for clarity. Gray lines connect paired observations from the same participant, illustrating individual response to different montage conditions. Large colored circles represent group means with vertical error bars indicating +-1 standard deviation. Statistical comparisons were performed using paired t-test of Wilcoxon signed-rank test with significance denoted as: p < 0.001, p < 0.01, p < 0.05, n.s. = not significant. Effect size (Cohen’s d/r) and percentage change are displayed above each comparison. TI_max_ are expressed as V/m.

The right hippocampus demonstrated comparable patterns with mean TI_max_ values decreased from 0.316 V/m in optimized solutions to 0.305 V/m when mapped (Δ = -0.0112 V/m, -3.67%, p = 1.07×10⁻⁷), while maximum values decreased from 0.510 V/m to 0.495 V/m (Δ = -0.0147 V/m, -2.98%, p = 8.22×10⁻⁷). Focality also showed a small but statistically significant reduction from 1.526 to 1.487 (-2.62%, p = 0.000642), indicating modest compromise in spatial selectivity for this deeper target (Fig 5 C).

For the spherical target, mapping effects were intermediate between the two anatomical targets. Mean TI_max_ decreased from 0.350 V/m to 0.342 V/m (Δ = -0.0079 V/m, -2.31%, p = 2.66×10⁻⁵), with maximum values showing similar reductions from 0.386 V/m to 0.377 V/m (Δ = -0.009 V/m, -2.33%, p = 6.66×10⁻⁵). Focality remained statistically unchanged (1.547 vs 1.555, p = 0.263), reinforcing that electrode mapping preserves targeting precision (Fig 5 B).

### 3.3 Individualized versus generalized head models

We next examined whether using individualized head models provides meaningful advantages over a generalized template (Ernie model, sub37) for TI optimization. To avoid self-comparison, the generalized model subject was excluded from these analyses (n = 36). For the left insula, individualized models with mapped electrodes achieved higher field intensities than the generalized model, with mean TI_max_ of 0.321 V/m versus 0.312 V/m (+2.70%, p = 0.0002) and maximum values of 0.464 V/m versus 0.450 V/m (Δ = +0.015 V/m, +3.31%, p = 0.0009). Focality showed no significant difference between approaches (1.418 vs 1.418, p = 0.990) (Fig 6 A, D).

**Figure 6.**
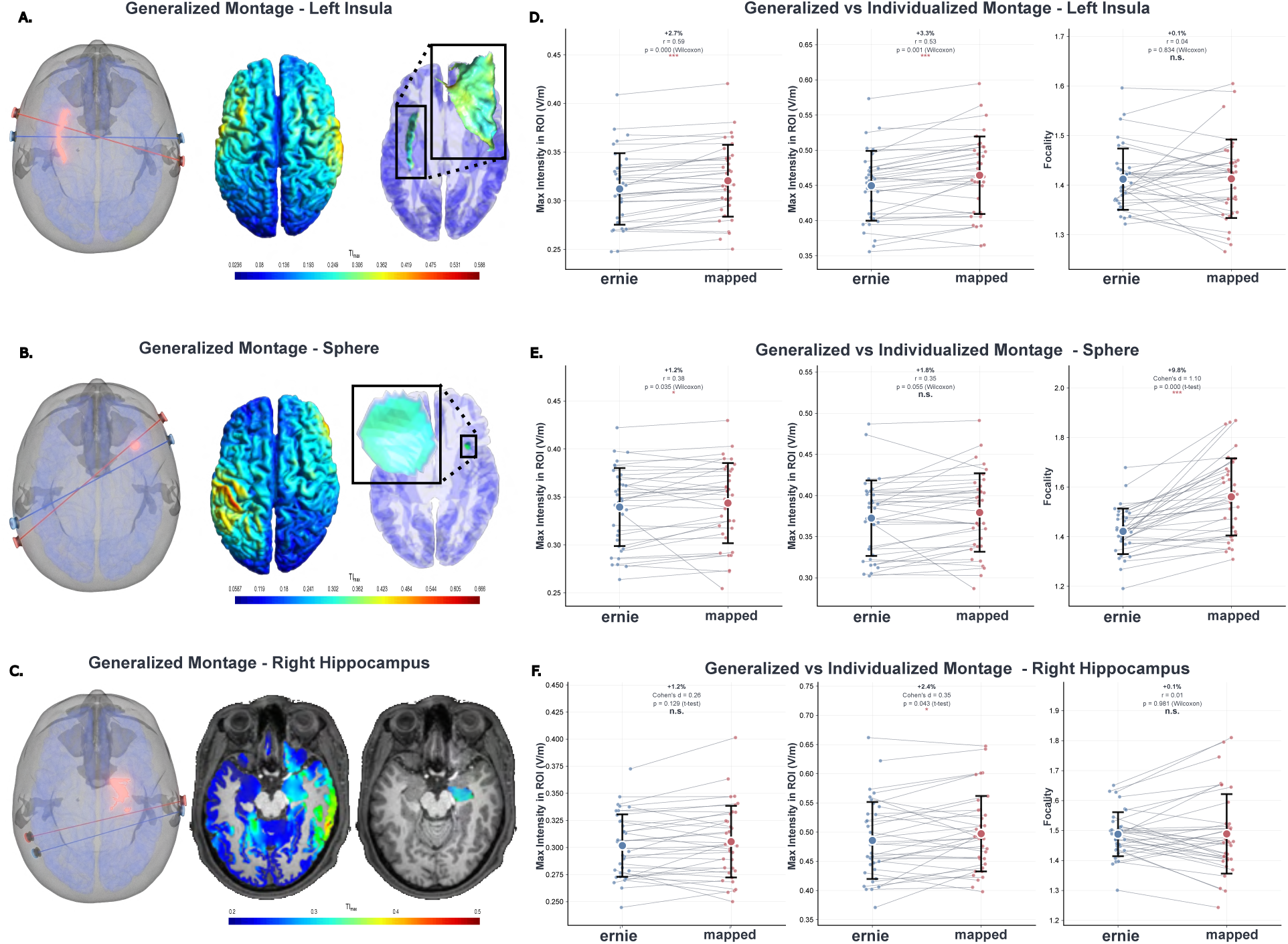
Individualized versus generalized head modeling performance. **A-C.** On the left, montages suggested by the generalized model (subject 37, ERNIE) for the Left insula, sphere, and right hippocampus. On the right, cortical field distribution and extracted ROI for subsequent analyses. **D.** Left Insula comparison: subject-specific suggested mapped montages (red dots) versus generalized template montage (ernie, blue dots). Individualized models achieved higher mean TI_max_ (+2.70%, p = 0.0002) and maximum TI_max_ (+3.31%, p = 0.0009) compared to template. **E.** Spherical comparison: Individualized models showed modest advantages in maximum field (+1.2%, p = 0.035) and superior focality (+9.8%, p = 8.91×10⁻⁶). **F.** Right hippocampus comparison in volumetric space: Comparable field intensities between approaches (mean TImax: +1.21%, p = 0.129; maximum TImax: +2.38%, p = 0.0433) with no significant difference in focality (+0.07%, p = 0.981). Individual participants are shown as small colored dots (n=36, excluding the generalized model subject) with slight horizontal jitter for clarity. Gray lines connect paired observations from the same participant, illustrating individual response to different montage conditions. Large colored circles represent group means with vertical error bars indicating +-1 standard deviation. Statistical comparisons were performed using paired t-test or Wilcoxon signed-rank test with significance denoted as: p < 0.001, p < 0.01, p < 0.05, n.s. = not significant. Effect size (Cohen’s d/r) and percentage change are displayed above each comparison. TImax are expressed as V/m.

**Figure 7.**
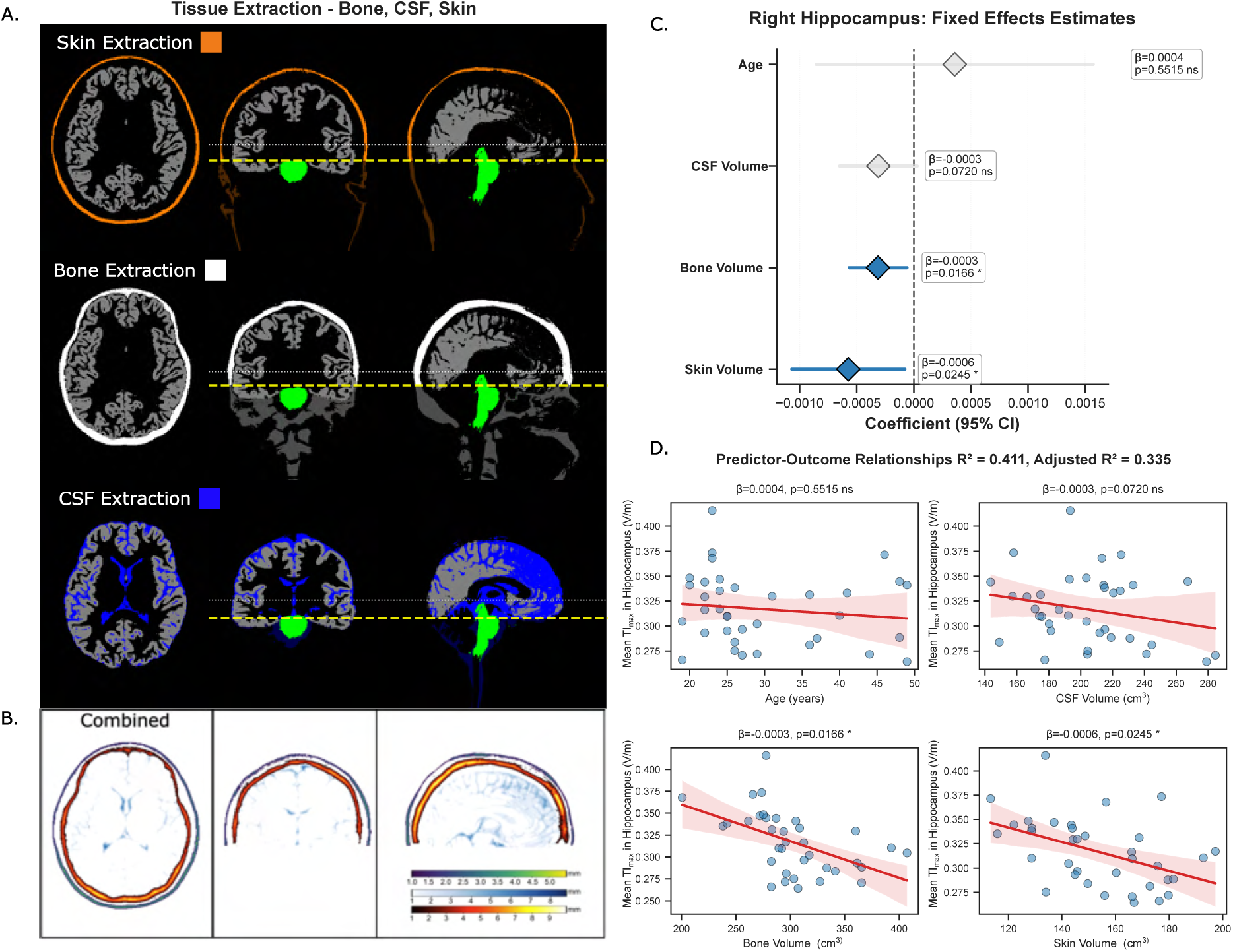
Anatomical determinants of inter-individual variability in TI field exposure. **A.** Representative tissue segmentation from a single participant displayed in axial, coronal, and sagittal views. Skin (orange), gray matter (grey) and cortical bone structures (white, including compact and spongy bone layers), cerebrospinal fluid (CSF) highlighting subarachnoid and ventricular CSF distributions (dark blue). **B.** Visualization of skin, skull and CSF mean thickness mapping, with color gradient indicating thickness variations (scale: X-Y mm). **C.** Multiple linear regression coefficient forest plot for right hippocampus showing standardized coefficients and 95% confidence intervals. Bone volume (β = -0.000314, p = 0.0166) and skin volume (β = -0.000574, p = 0.0245) emerged as significant predictors, CSF volume approached significance (β = -0.000311, p = 0.0720), while age showed no association (β = 0.000358, p = 0.5515). **D.** Scatter plots demonstrating predictor-outcome relationships for right hippocampus TI_max_ field intensity. Individual data points represent participants (n=36) with regression lines and significance indicators (*p<0.05, n.s. = not significant).

The right hippocampus revealed an interesting pattern where field intensity metrics were comparable between individualized and generalized models. Mean TI_max_ values were virtually identical (0.305 vs 0.302 V/m, Δ = +0.00365 V/m, +1.21%, p = 0.129), while maximum values showed a modest significant increase (0.497 vs 0.486 V/m, Δ = +0.0116 V/m, +2.38%, p = 0.0433). Focality showed no significant difference between approaches (1.489 vs 1.488, +0.07%, p = 0.981), suggesting comparable spatial selectivity for this deep target (Fig6 C, F).

For the spherical target, individualized models showed modest advantages in maximum field values (0.377 vs 0.371 V/m, Δ = +0.006 V/m, +1.62%, p = 0.0411) but not mean values (0.342 vs 0.338 V/m, Δ = +0.004 V/m, +1.18%, p = 0.0638). Notably, individualized models achieved substantially better focality compared to the template (1.555 vs 1.420, +9.53%, p = 8.91×10⁻⁷), indicating superior spatial selectivity when targeting small regions (Fig6 B, E).

### 3.4 Determinants of inter-individual variability

To identify anatomical and demographic factors underlying inter-individual variability in TI field exposure, we analyzed data from the optimized electrode montages for both the left insula and right hippocampus. Initial correlation analysis revealed strong within-tissue correlations (Figure S4), with bone thickness and volume showing correlations exceeding r = 0.9, and similar patterns for CSF and skin measures. Based on this multicollinearity assessment, we selected volume measures over thickness measures for subsequent analyses.

Multiple Linear Regression Analysis: To quantify the overall variance explained by anatomical factors, we performed MLR analyses with age, CSF volume, bone volume, and skin volume as predictors of TI field intensity. For the left insula, these anatomical features collectively explained 46.9% of the variance in field intensity (R² = 0.469, adjusted R² = 0.400, F = 6.833, p = 4.59×10⁻⁴), indicating substantial predictive power. Individual predictors showed that bone volume (β = -0.000430, p = 0.0024) and skin volume (β = - 0.000600, p = 0.0248) were significant predictors of field intensity, while CSF volume (β = - 0.000113, p = 0.5227) and age (β = -0.000134, p = 0.8315) showed no significant associations.

The right hippocampus model showed comparable explanatory power, with anatomical features accounting for 41.1% of variance (R² = 0.411, adjusted R² = 0.335, F = 5.411, p = 2.01×10⁻³). Similarly, bone volume (β = -0.000314, p = 0.0166) and skin volume (β = - 0.000574, p = 0.0245) emerged as significant predictors. CSF volume showed a marginally significant association (β = -0.000311, p = 0.0720), approaching the significance threshold, while age remained non-significant (β = 0.000358, p = 0.5515). These findings confirm that tissue-specific anatomical features, particularly bone and skin volumes, are robust predictors of TI field penetration across different brain targets.

### 3.5 Surface-aligned field component

Given the importance of field orientation relative to cortical geometry (12,46,47), we analyzed the normal component of TI fields (TI_normal_) for cortical targets. Comparing mapped to optimized montages for the left insula, mean TI_normal_ decreased from 0.226 V/m to 0.216 V/m (Δ = -0.0096 V/m, -4.43%, p = 2.96×10⁻⁹), with maximum values showing similar reductions from 0.402 V/m to 0.394 V/m (Δ = -0.008 V/m, -1.99%, p = 0.0157).

Normal focality remained statistically unchanged (2.265 vs 2.244, p = 0.117), suggesting preserved directional selectivity despite electrode constraints (Fig 8A). For the spherical target (using the max optimization that maximized TI_normal_), comparable sensitivity of the normal component to electrode mapping was observed, with mean values decreasing from 0.201 V/m to 0.192 V/m (Δ = -0.0094 V/m, -4.89%, p = 5.22×10⁻⁶) and maximum values from 0.351 V/m to 0.343 V/m (Δ = -0.008 V/m, -2.28%, p = 0.00412). Normal focality showed a trend toward reduction that did not reach significance (2.025 vs 1.983, p = 0.0681) (Fig 8C).

**Figure 8.**
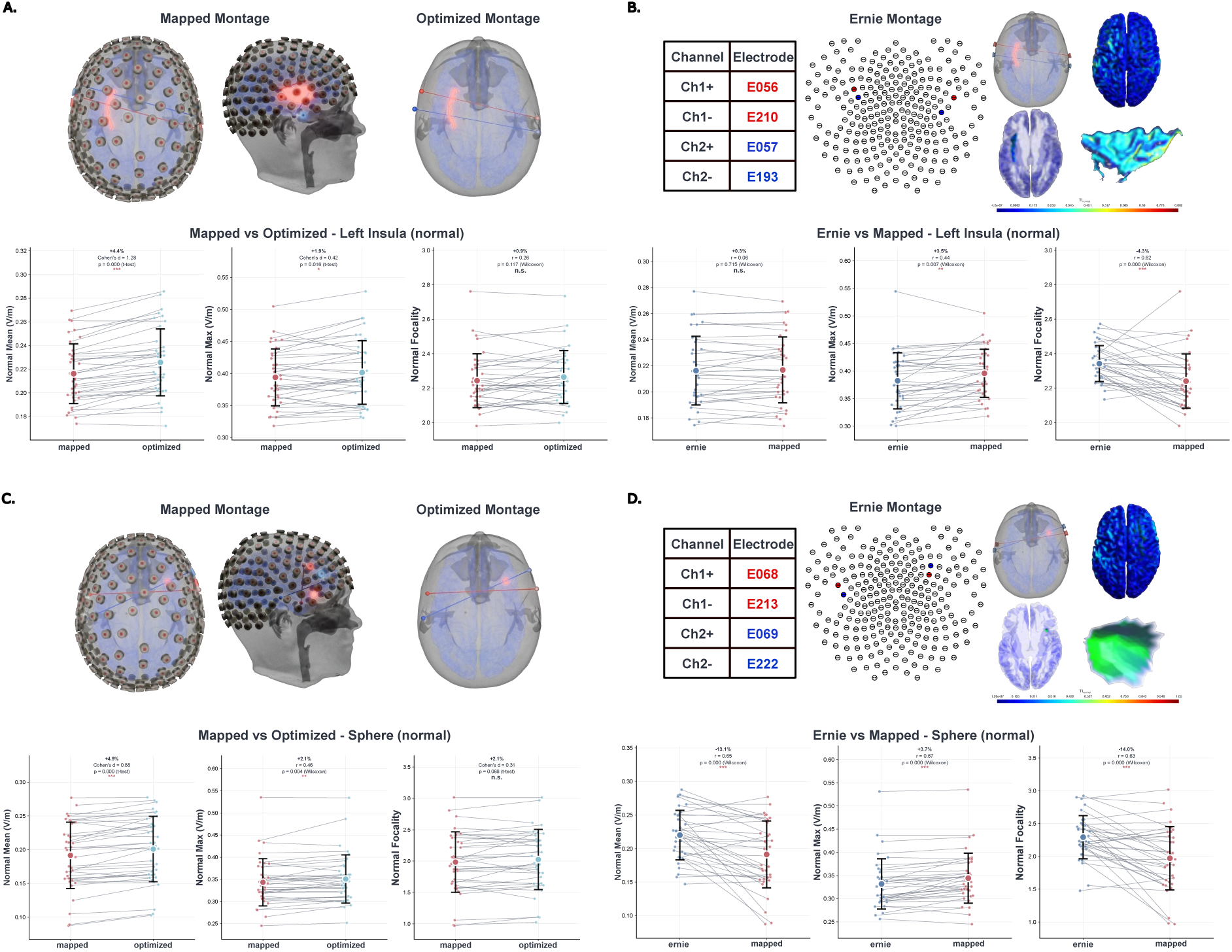
Surface-aligned normal component (TI_normal_) analysis for cortical and spherical targets. **A.** Left insula comparison between optimized and mapped electrode configurations for TI_normal_ field component (optimization goal: maximize mean TI_normal_). Mean values decreased from 0.226 to 0.216 V/m (Δ = -4.43%, p = 2.96×10⁻⁹), while focality remained preserved (p = 0.117). **B.** Left insula comparison between individualized mapped montages and generalized model (Ernie) for TI_normal_ (optimization goal: maximize mean TI_normal_). No significant difference in mean values (0.216 V/m for both, p = 0.905), though individualized models achieved higher maximum values (+3.41%, p = 0.0181) with reduced focality (-4.20%, p = 0.000112). **C.** Spherical target comparison between optimized and mapped configurations for TI_normal_ (optimization goal: maximize maximum TI_normal_). Mean values decreased from 0.201 to 0.192 V/m (Δ = -4.89%, p = 5.22×10⁻⁶), with non-significant focality reduction (p = 0.0681). **D.** Spherical target comparison between individualized and generalized models for TI_normal_ (optimization goal: maximize maximum TI_normal_). Individualized models showed lower mean values (0.192 vs 0.220 V/m, -12.75%, p = 0.000153) but higher maximum values (+3.65%, p = 0.000175) with substantially reduced focality (-13.61%, p = 0.000302). All panels display individual participant data points with paired connections, group means ± standard deviation, and statistical comparisons using paired t-test or Wilcoxon signed-rank test. Significance levels: ***p < 0.001, **p < 0.01, *p < 0.05, n.s. = not significant.

When comparing individualized mapped montages to the generalized model, the left insula showed no difference in mean TI_normal_ (0.217 V/m for mapped vs 0.216 V/m for ernie, p = 0.715), though individualized models achieved higher maximum values (0.396 vs 0.382 V/m, Δ = +0.013 V/m, +3.52%, p = 0.00738) at the cost of reduced focality (2.241 vs 2.342, - 4.32%, p = 8.47×10⁻⁵) (Fig 8B). The spherical target demonstrated more pronounced differences, with individualized models showing lower mean TI_normal_ (0.191 vs 0.220 V/m, - 13.08%, p = 0.000145) but higher maximum values (0.344 vs 0.332 V/m, +3.75%, p = 0.000175) and substantially reduced focality (1.971 vs 2.293, -14.01%, p = 0.000295) (Fig8 D). These results suggest that while the normal component provides additional insight into field-tissue interactions, it does not demonstrate superior sensitivity to methodological choices compared to TI_max_.

## 4. Discussion

### 4.1 TI-Toolbox: strengths and the gap it closes

TI-Toolbox unifies the end-to-end TI workflow—from DICOM preprocessing and head- model generation through montage optimization, field simulation, analysis, and visualization —under a single, containerized framework with both GUI and CLI access. The platform lowers technical barriers, promotes standardized outputs and reports, and enables reproducible, cohort-level studies. Beyond orchestration, it adds TI-specific capabilities: electrode mapping from free solutions to standard HD-EEG nets, support for multiple optimization approaches, computation of both TI_max_ and grey matter surface- aligned TI_normal_, and multi-polar simulations.

### 4.2 Related software and the SimNIBS foundation

Several existing tools provide components for electrical stimulation modeling, each with distinct strengths and limitations. Open-source platforms such as ROAST (48) and COMETS2 (49) offer valuable functionality for conventional tES applications but lack native TI support and require extensive customization for multi-stage TI workflows. Commercial solutions like the TI Planning Tool (50) and HD-Targets-IFS (51) provide advanced TI-specific features but remain closed-source and can be costly or operationally constrained for many research groups.

We chose to build TI-Toolbox as an integrated wrapper around SimNIBS (52) for several compelling reasons. First, SimNIBS provides a Python-based interface that enables programmatic access to all core functionalities, facilitating easy integration. This allowed us to extend its capabilities while maintaining compatibility with its core functionalities. Second, SimNIBS has been validated through peer-reviewed studies comparing simulated fields with intracranial recordings (14,53). Lastly, SimNIBS benefits from an active community of developers and users who continuously contribute improvements, bug fixes, and methodological advances.

### 4.3 Case studies: summary and interpretation

Our case studies reveal several important insights for the practical implementation of TI stimulation. While many of our comparisons reached statistical significance, it is crucial to distinguish between statistical and clinical relevance. The cohort-level mean percentage differences in ROI intensity were typically small, ranging from approximately 1–4%, and in isolation are unlikely to be clinically meaningful as intensity remains far within sub- threshold stimulation regime (54–56). However, dedicated efforts should investigate the possibility of using TI for neural entrainment where such field variance may be critical (57,58).

The validity of electrode mapping emerged as a particularly encouraging finding for clinical translation. When we mapped unconstrained genetic optimization solutions to standard HD-EEG net positions, the resulting montages preserved both intensity and focality characteristics without substantially impairing optimization benefits. This preservation of field characteristics suggests that mapping offers a practical route to montage standardization and smoother clinical translation, eliminating the need for specialized neuronavigation devices while maintaining targeting efficacy. The successful mapping validates the use of pre-manufactured HD-EEG nets in clinical settings, potentially reducing both cost and complexity of TI implementation.

A comparison between individualized and generalized head models revealed a nuanced picture of when each approach is most appropriate. The generalized head model proved adequate for discovering effective electrode configurations for both cortical and subcortical targets, suggesting that initial montage design and optimization can proceed without individual MRI scans. However, accurate exposure assessment clearly requires individualized models due to the substantial inter-individual variability observed in our cohort in line with previous research (2,21,59). This finding supports a hybrid workflow where generalized models guide initial protocol development, while individualized models enable precise dose determination and safety assessment for clinical applications (16,17). It is important to note that our analysis assumes quasi-uniform field distributions within ROIs when computing mean values, while in reality the fields exhibit spatial gradients and hotspots that may have differential physiological effects depending on the specific neural circuits engaged (60).

The analysis of inter-individual variability revealed that tissue composition plays a critical role in determining field exposure patterns. Consistent with previous tES literature(61–63), anatomical features of the head substantially influence electric field penetration and distribution. Our multiple linear regression analyses quantified these relationships, demonstrating that anatomical features (age, CSF volume, bone volume, and skin volume) collectively explained 46.9% of variance in field intensity for the left insula and 41.1% for the right hippocampus using optimized montages. Bone volume and skin volume emerged as significant predictors for both targets, confirming their consistent role as barriers to electric field penetration. Interestingly, CSF volume showed a marginally significant association (p = 0.0720) for the right hippocampus but not for the left insula (p = 0.5227), likely reflecting the hippocampus’s proximity to the ventricular system where CSF-filled spaces create more pronounced current shunting effects. Age showed no significant association with field intensity in either target, suggesting that anatomical characteristics mediate any age-related effects on field penetration.

Finally, our evaluation of surface-aligned field components addressed an important consideration for cortical stimulation. Traditionally, the normal direction of an exogenous electric field has been proposed to be critical, due to the orientation of pyramidal neurons perpendicular to the cortical surface (64,65). Our analyses revealed that TI_normal_ did not demonstrate superior sensitivity to methodological manipulations compared to TI_max_ when evaluating mapping effects or model individualization. Nevertheless, we recommend reporting both metrics for completeness, as the physiological relevance of field directionality remains an active area of investigation in the TI literature (12).

### 4.4 Limitations

Several limitations warrant consideration when interpreting our findings. First, our simulations employed isotropic conductivity values and did not leverage diffusion-derived anisotropy which could influence field exposure. However, there is not enough evidence that an anisotropic approach is necessary for targeting gray matter ROIs (55). Future research should systematically investigate the impact of incorporating DTI on TI field distributions.

Second, all electromagnetic field calculations relied on the quasi-static approximation, which assumes that propagation and inductive effects are negligible at the frequencies employed in TI stimulation (typically 1-20 kHz carriers) (3). While this approximation is widely accepted for conventional tES and has been validated for frequencies up to several kilohertz in biological tissues (66), the validity at TI carrier frequencies warrants continued investigation (67).

Third, our electrode mapping approach, while successful in preserving field characteristics, was evaluated using only the inner 185 electrodes of the GSN-HydroCel- 256 system (EGI/Philips) (68). This high-density array provides smaller inter-electrode spacing compared to traditional EEG nets, which may represent an upper bound for successful mapping resolution. The minimum electrode density required for accurate mapping of optimized TI montages remains unexplored, and future work should systematically evaluate how mapping fidelity degrades with decreasing electrode density, particularly for standard 10-20 with larger inter-electrode distances.

Finally, the tissue analyzer module represents an experimental tool that, while providing consistent measurements of anatomical properties across our cohort, has not been independently validated against ground-truth measurements from high-resolution CT imaging (69) or other specialized segmentation methods (70). Nevertheless, it serves as an informative tool for researchers performing computational modeling by automatically extracting key anatomical metrics that our results—and the broader tES literature— consistently identify as primary contributors to inter-individual variability. The automated extraction of tissue metrics provides researchers with readily available covariates that can explain substantial portions of field variance. This functionality aims to support hypothesis generation and covariate adjustment in group studies rather than provide clinical-grade morphometric assessment.

### 4.5 Implications and future directions

The development and validation of TI-Toolbox carries several implications for the broader TI research community. As an open-source platform, the toolbox provides an opportunity for collaborative advancement of TI methodologies. We encourage researchers to not only utilize the platform but also to contribute improvements and share workflows in its dedicated GitHub Discussion page (71). This community-driven approach will be essential for establishing standardized protocols and accelerating the development of interoperable tools across different research groups and clinical centers.

Our findings also inform best practices for clinical implementation of TI stimulation. The evidence suggests that prospective clinical studies should adopt a tiered approach to model complexity. While generalized head models can effectively guide initial montage design and protocol development, individualized models become crucial when relating exposure to electrophysiological and behavioral outcomes (72,73). This distinction is particularly important for dose-response studies and for understanding inter-individual variability in treatment response (74). The substantial variance in field exposure across individuals, even with identical montages, underscores the necessity of subject-specific modeling for precision medicine applications.

Regarding field assessment metrics, our analyses support reporting strategies that capture multiple aspects of TI stimulation. We recommend that researchers report both TI_max_ and TI_normal_ for cortical targets, as these metrics provide complementary information about field intensity and directionality. While TI_max_ remains the primary metric for overall exposure assessment, TI_normal_ may prove particularly relevant for understanding interactions with columnar cortical organization (64,65).

Furthermore, we recommend optimizing for mean field values within the ROI rather than maximum values. Our analyses revealed that when optimization targeted mean TI_normal_, the maximum values also increased correspondingly, but the reverse was not true (Figure 8).

This approach promotes more homogeneous field distributions across the target region, potentially engaging a larger proportion of the neural population while avoiding hotspots that may dominate max-based optimization strategies.

Finally, our experience with optimization algorithms provides guidance for practical implementation. The flex-search algorithm achieves field intensities comparable to, if not better than, existing optimization algorithms reported in the literature (75). The multi-start strategy with three independent runs effectively minimizes the optimization function value and improves targeting, as demonstrated for the spherical ROI where it yielded a 4.18% improvement (p = 5.48×10⁻⁶). However, this modest gain must be weighed against the tripled computational cost, suggesting that single runs may suffice for many applications (Figure S1). Validation of flex-search results using ex-search, where we evaluated four candidate electrode positions around each optimized electrode location, agreed with the flex-search suggestions, confirming the robustness of the genetic algorithm and mapping approach (Table S2). The focality optimization objective proved more nuanced and context-dependent, requiring careful consideration of thresholding strategies. Our analysis revealed that dynamic thresholding dramatically affects results - 50% relative thresholds increased focality by 75% compared to fixed thresholds, while 80% thresholds decreased it by 36.6%. This suggests that optimal threshold selection is critical for successful optimization (Figure S3) (32). Further research should be conducted into the relationship of the dynamic thresholding and ROI location as it is possible that more superficial ROIs will benefit from high upper bound thresholds while converging successfully on a more focal solution.

### 4.6 Concluding remarks

Taken together, the case studies support a pragmatic translational pathway: use standardized mapping for clinical usability, leverage a generalized model to prototype montages, and rely on individualized models for exposure quantification and safety assessment. TI-Toolbox operationalizes this pathway within a reproducible, open, and extensible framework, laying the groundwork for multi-center harmonization and prospective validation.

## Acknowledgements

We wish to thank the staff at University of Wisconsin Brain Imaging Waisman Center and Center for Healthy Minds where the MRI data was collected. We appreciate all study participants for their time and contribution to this work.

## CReDIT authorship contribution statement

**Ido Haber:** Writing – review & editing, Writing – original draft, Visualization, Software, Methodology, Formal analysis, Data curation, Conceptualization. **Aksel Jackson**: Writing – review & editing, Writing – original draft, Visualization, Software. **Axel Thielscher**: Writing – review & editing, Software, Methodology. **Aviad Hai**: Writing – review & editing. **Giulio Tononi**: Writing – review & editing, Conceptualization, Funding acquisition.

## Data Availability

- Code repository: https://github.com/idossha/TI-Toolbox

- Documentation and tutorials: https://idossha.github.io/TI-Toolbox/

## Declaration of generative AI use

During the preparation of this work, the author(s) used Cursor for text and code editing. After using this tool/service, the author(s) reviewed and edited the content as needed and take(s) full responsibility for the content of the published article.

