## Supplemental Information for "TI-Toolbox: An Open-Source Software for Temporal Interference Stimulation Research"

*Supplementary Information*

List of contents


| | | |
|---|---|---|
| Table S1. | Mapping of optimized electrode to HD-EEG net | 2 |
| Table S2. | Exhaustive search evaluation | 3 |
| Table S3 | Isotropic conductivity values | 4 |
| Table S4 | Demographic and anatomical descriptive information | 4 |
| Figure S1 | Multi-start optimization | 5 |
| Figure S2 | Multipolar vs Unipolar simulation Targeting | 6 |
| Figure S3 | Sphere focality under fixed and relative thresholds | 7 |
| Figure S4 | Correlation matrix of anatomical and demographic predictors | 8 |
| Figure S5 | Mapping of optimized electrode statistics | 9 |

The transition from unconstrained optimization solutions to practical electrode montages represents a critical step in clinical translation. While genetic algorithms can identify theoretically optimal electrode positions anywhere on the scalp, its transition to clinical application may be difficult. Our electrode mapping algorithm bridges this gap by finding the best approximation of optimized positions using available electrode sites. For this study, we utilized the inner 185 electrodes of the GSN-HydroCel-256 system (EGI/Philips), which provides high-density coverage. This HD-EEG configuration was chosen for several reasons: (1) it offers sufficient spatial resolution to approximate most optimized positions within clinically acceptable distances, (2) the electrodes cover the majority of the scalp surface accessible for stimulation, excluding neck regions and face, and (3) it represents a commercially available system already in use. The mapping process employs the Hungarian algorithm, a combinatorial optimization method that solves the assignment problem in polynomial time. By minimizing the total Euclidean distance between optimized and standard positions, this approach ensures good representation of the intended field distribution while maintaining practical feasibility. Tables S1 demonstrates representative mapping results, showing that the mean mapping distances typically range from 4-9 mm for the insula. For a deeper investigation to the different regions analyzed see Figure S5.

| Electrode | Mapped to | Original position | Mapped position | Distance |
|---|---|---|---|---|
| 0 (Channel 0, Array 0) | E063 | [-77.08, 28.24, 22.85] | [-77.68, 24.18, 24.69] | 4.49 |
| 1 (Channel 0, Array 1) | E203 | [82.25, 17.16, 14.56] | [81.80, 22.65, 22.60] | 9.75 |
| 2 (Channel 1, Array 0) | E193 | [82.97, -5.17, 23.04] | [83.29, 0.92, 18.94] | 7.35 |
| 3 (Channel 1, Array 1) | E056 | [-73.98, 37.68, 28.71] | [-71.69, 45.56, 28.90] | 8.21 |

**Table S1.** Mapping of optimized electrode to HD-EEG net for the left insula (ernie). The table shows the original optimized electrode coordinates, their mapped positions on the HD-EEG net, and the Euclidean distance between original and mapped positions. Channel 0 and Channel 1 represent the two TI channel pairs, each with two electrodes (Array 0 and Array 1).

Following the mapping of optimized electrode positions to standard EEG locations, we validated our genetic algorithm results through an exhaustive search of nearby electrode combinations. This validation step serves two purposes: confirming that the flex-search algorithm identified near-optimal solutions and exploring whether small adjustments to mapped positions could yield meaningful improvements. The exhaustive search systematically evaluated electrode configurations in the neighborhood of the mapped solution:

**E1+:** E001, E002, E003, E010 || **E1-:** E076, E077, E086, E087
**E2+:** E059, E066, E060, E052 || **E2-:** E222, E221, E220, E212

This resulted in 256 unique combinations ($4^4$ configurations). The top-performing configuration (E001_E077 <> E060_E221) achieved a mean $TI_{max}$ of 0.285 V/m within the ROI, remarkably close to the flex-search result of 0.284 V/m. This near-perfect agreement (0.35% difference) provides strong validation that our genetic algorithm effectively navigates the complex optimization landscape, finding solutions that are robust to small perturbations in electrode positioning.

| Electrode Configuration | Max $TI_{max}$ ROI | Mean $TI_{max}$ ROI | Peak Field | 95th % | 99th % | 99.9th % | Focality 50% | Focality 75% | Focality 90% | Focality 95% | Peak Location |
|---|---|---|---|---|---|---|---|---|---|---|---|
| E001_E077 <> E060_E221 | 0.293 | 0.285 | 1.202 | 0.313 | 0.4 | 0.567 | 70002 | 5276.9 | 1697.3 | 1168.5 | 16.54, 51.14, -5.46 |
| E001_E077 <> E066_E221 | 0.293 | 0.284 | 1.218 | 0.334 | 0.423 | 0.573 | 82545.3 | 7036.9 | 1777.8 | 1166.7 | 16.54, 51.14, -5.46 |
| E001_E077 <> E052_E221 | 0.293 | 0.284 | 1.249 | 0.297 | 0.39 | 0.569 | 50784.9 | 4532.3 | 1668.4 | 1172.9 | 16.54, 51.14, -5.46 |
| E001_E077 <> E059_E221 | 0.293 | 0.284 | 1.264 | 0.313 | 0.406 | 0.574 | 63095.9 | 5308 | 1674.9 | 1150.3 | 16.54, 51.14, -5.46 |
| E001_E087 <> E060_E221 | 0.293 | 0.284 | 1.202 | 0.305 | 0.39 | 0.566 | 65266.5 | 4460.8 | 1611.4 | 1145.6 | 16.54, 51.14, -5.46 |
| E001_E087 <> E052_E221 | 0.293 | 0.284 | 1.249 | 0.292 | 0.385 | 0.569 | 46263.4 | 4294.3 | 1663.4 | 1176.6 | 16.54, 51.14, -5.46 |
| E002_E077 <> E066_E221 | 0.292 | 0.284 | 1.097 | 0.337 | 0.426 | 0.571 | 89265.6 | 7546.3 | 1853.2 | 1201.1 | 16.54, 51.14, -5.46 |
| E002_E087 <> E066_E221 | 0.29 | 0.284 | 1.047 | 0.319 | 0.397 | 0.565 | 81010.5 | 4889.6 | 1659.1 | 1163.9 | 16.54, 51.14, -5.46 |
| E001_E087 <> E066_E221 | 0.292 | 0.283 | 1.218 | 0.316 | 0.396 | 0.57 | 73254.6 | 4592.5 | 1601.3 | 1120.9 | 16.54, 51.14, -5.46 |
| E001_E087 <> E059_E221 | 0.292 | 0.283 | 1.264 | 0.299 | 0.391 | 0.573 | 51689.7 | 4295.8 | 1617.8 | 1140.3 | 16.54,51.14,-5.46 |
| E003_E076 <> E059_E220 | 0.251 | 0.242 | 0.983 | 0.285 | 0.357 | 0.451 | 155009.5 | 11964.8 | 2626.9 | 1492.3 | 25.85, 25.24, -19.94 |
| E003_E087 <> E059_E220 | 0.25 | 0.242 | 0.856 | 0.274 | 0.333 | 0.43 | 182103.3 | 10190.7 | 2475.6 | 1491.9 | 25.85, 25.24, -19.94 |
| E003_E086 <> E059_E220 | 0.25 | 0.242 | 0.939 | 0.275 | 0.335 | 0.427 | 191006.9 | 11590.7 | 2509.9 | 1517 | 25.85, 25.24, -19.94 |
| E003_E076 <> E060_E220 | 0.25 | 0.242 | 0.983 | 0.283 | 0.347 | 0.445 | 161179.9 | 10953.4 | 2389.3 | 1420.8 | 25.85, 25.24, -19.94 |
| E003_E087 <> E060_E220 | 0.249 | 0.242 | 0.856 | 0.285 | 0.347 | 0.44 | 189361.7 | 11949.2 | 2660.8 | 1477.7 | 25.85, 25.24, -19.94 |
| E003_E086 <> E060_E220 | 0.249 | 0.242 | 0.939 | 0.28 | 0.339 | 0.429 | 197434.1 | 12275.8 | 2507.8 | 1500 | 25.85, 25.24, -19.94 |
| E003_E077 <> E052_E220 | 0.249 | 0.241 | 0.919 | 0.276 | 0.337 | 0.432 | 177965 | 10890.3 | 2307.6 | 1412.1 | 25.85, 25.24, -19.94 |
| E003_E076 <> E052_E220 | 0.249 | 0.241 | 0.983 | 0.27 | 0.329 | 0.424 | 181214.5 | 10358.5 | 2136.4 | 1363.5 | 25.85, 25.24, -19.94 |
| E003_E087 <> E052_E220 | 0.248 | 0.241 | 0.856 | 0.267 | 0.321 | 0.416 | 205468.8 | 10436.8 | 2238.1 | 1370.4 | 25.85, 25.24, -19.94 |
| E003_E086 <> E052_E220 | 0.248 | 0.24 | 0.939 | 0.265 | 0.32 | 0.415 | 196756.6 | 9883.6 | 2130.4 | 1315.2 | 25.85, 25.24, -19.94 |

**Table S2.** Example output from the ex-search optimization process. Results from exhaustive search (ex-search) algorithm based on the Ernie mapped electrode configuration targeting the spherical ROI (MNI: 36.10, 14.14, 0.33; radius 5 mm) while maximizing mean $TI_{max}$ (highlighted in yellow). The table shows the top 10 and bottom 10 electrode combinations separated by the red line (out of 256 combinations) ranked by mean $TI_{max}$ intensity within the ROI, including focality metrics at various percentiles and peak field locations.

All simulations employed isotropic conductivity values based on established literature. While anisotropic conductivity modeling is available in TI-Toolbox through SimNIBS's DTI integration, the isotropic approach was used for this study.

| Tissue Number | Tissue Name | Conductivity (S/m) |
|---|---|---|
| 1 | White matter | 0.126 |
| 2 | Gray matter | 0.275 |
| 3 | Cerebrospinal fluid (CSF) | 1.654 |
| 4 | Bone | 0.01 |
| 5 | Scalp | 0.465 |
| 6 | Eyeballs | 0.5 |
| 7 | Compact Bone | 0.008 |
| 8 | Spongy Bone | 0.025 |
| 9 | Blood | 0.600 |
| 10 | Muscle | 0.160 |
| 100 | Silicone | 29.4 |
| 500 | Saline | 1.0 |

**Table S3**. Isotropic tissue conductivity values employed in all TI simulations. These values represent the electrical conductivity (σ) of biological tissues at the kilohertz frequencies typical of TI stimulation (1-20 kHz). The conductivity hierarchy—from highly conductive CSF (1.654 S/m) and blood (0.6 S/m) to resistive bone (0.008-0.025 S/m)—fundamentally shapes current flow patterns through the head. Gray matter (0.275 S/m) and white matter (0.126 S/m) exhibit intermediate conductivities that influence field distributions within neural tissue. The stark conductivity contrast between CSF and bone (>200-fold difference) creates the primary challenge in transcranial stimulation: current preferentially flows through CSF-filled spaces rather than penetrating to deep brain targets. These standardized values, drawn from literature enable reproducible simulations while acknowledging that actual tissue conductivities vary with factors including age and pathology.

| **Age (yrs, mean ± std, range)** | 29.9 ± 9.7 (range 19-49) |
|---|---|
| **Sex (%female)** | 58.3% |
| **Cortical Bone Volume ($cm^3$)** | 302.83 ± 42.86 |
| **Mean Skull thickness (mm)** | 3.42 ± 0.36 |
| **Total CSF volume ($cm^3$)** | 203.61 ± 34.26 |
| **Mean CSF thickness (mm)** | 1.85 ± 0.2 |
| **Skin volume ($cm^3$)** | 153.28 ± 21.85 |
| **Mean skin thickness (mm)** | 1.823 ± 0.136 |

**Table S4**. Demographic & anatomical descriptive information of the 36 individuals. Ernie data was excluded as there was missing information regarding the demographics and acquisition sequence of its MRI data.

The multi-start strategy addresses a fundamental challenge in genetic algorithm optimization: the potential convergence to local maxima due to the stochastic nature of the search process. We investigated whether running multiple independent optimizations with different random seeds would yield superior solutions compared to single runs. This approach is particularly relevant for TI applications where small changes in electrode positions can significantly affect field distributions.

Our analysis compared single-run versus three-run optimization strategies across different brain targets, with the multi-start approach automatically selecting the best solution based on the objective function value.

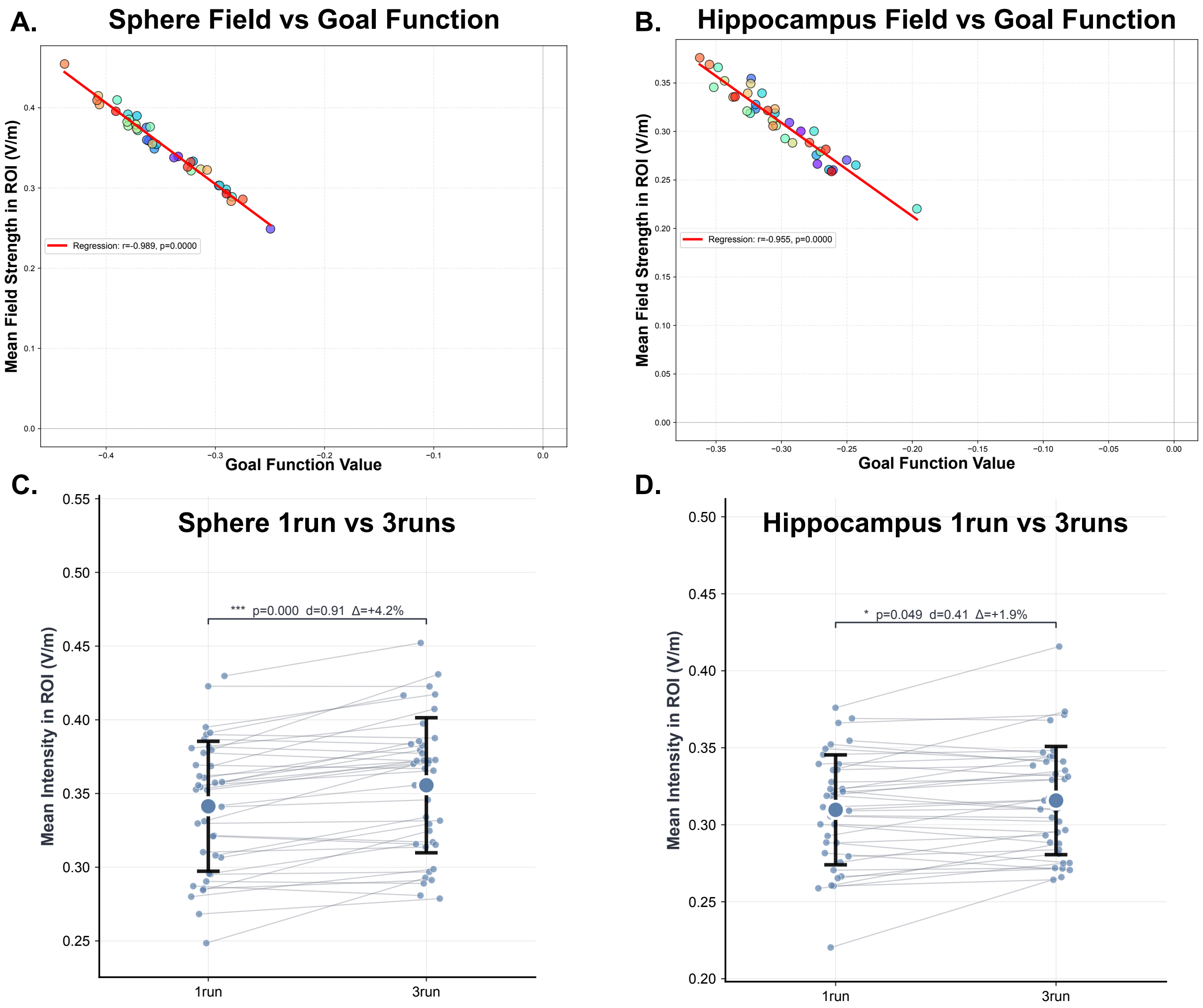


**Figure S1.** Multi-start optimization strategy evaluation. **A.** Relationship between optimization goal function values and achieved mean $TI_{max}$ field intensity in a 5 mm spherical ROI at MNI coordinates (36.10, 14.14, 0.33), demonstrating strong agreement between the flex-search algorithm's internal metric and post-hoc finite element method (FEM) simulations. Analysis performed in mesh space. **B.** Similar validation for right hippocampus target showing consistency between goal function optimization and simulated field outcomes in volumetric (voxel) space. **C.** Within-subject comparison of single-run versus three-run optimization for the spherical target. The multi-start approach yielded statistically significant improvement (mean $TI_{max}$: 0.355 vs 0.341 V/m; $\Delta = +0.014$ V/m, +4.18%, $p = 5.48\times10^{-6}$, Wilcoxon signed-rank test, $r = 0.683$, n=37). **D.** Hippocampus comparison showing modest but significant improvement with multi-start strategy (mean $TI_{max}$: 0.316 vs 0.310 V/m; $\Delta = +0.006$ V/m, +1.95%, $p = 0.049$, Wilcoxon signed-rank test, $r = 0.319$, n=37). Individual participant data points shown with paired connections, large colored circles represent group means ± standard deviation. While both targets show statistically significant improvements, the magnitude of enhancement is modest and should be weighed against the tripled computational cost.

Multi-polar temporal interference (mTI) represents an advanced stimulation paradigm that extends beyond conventional two-channel TI by employing multiple electrode pairs to achieve enhanced spatial control. By utilizing four or more independently controlled channels, mTI enables more degrees of freedom in field sculpting, potentially improving both focality and targeting flexibility. Our implementation demonstrates the computational framework required for mTI simulation, highlighting both its potential advantages and increased complexity. Multi-polar temporal interference (mTI) simulations were performed using the `simulator/mTI.py` module, which implements a hierarchical approach for combining multiple electrode pairs. The multi-polar configuration utilized four electrode pairs (8 electrodes total), labeled as HF_A, HF_B, HF_C, and HF_D. First, individual electric field simulations were computed for each electrode pair using SimNIBS's finite element method. Next, TI patterns were calculated pairwise: TI_AB was computed from the vector combination of HF_A and HF_B fields, and TI_CD from HF_C and HF_D fields, using the `get_TI_vectors` function that determines the modulation amplitude vectors based on field magnitudes and orientations. Finally, the multi-polar TI field (mTI) was calculated from the TI_AB and TI_CD using SimNIBS's `get_maxTI` function, which identifies the maximum modulation amplitude at each location. Both unipolar (2 electrode pairs) and multi-polar (4 electrode pairs) configurations used similar stimulation parameters: 2 mA total current injected (1mA per channel for uTI and 0.5mA for mTI per channel), 8 mm diameter electrodes.

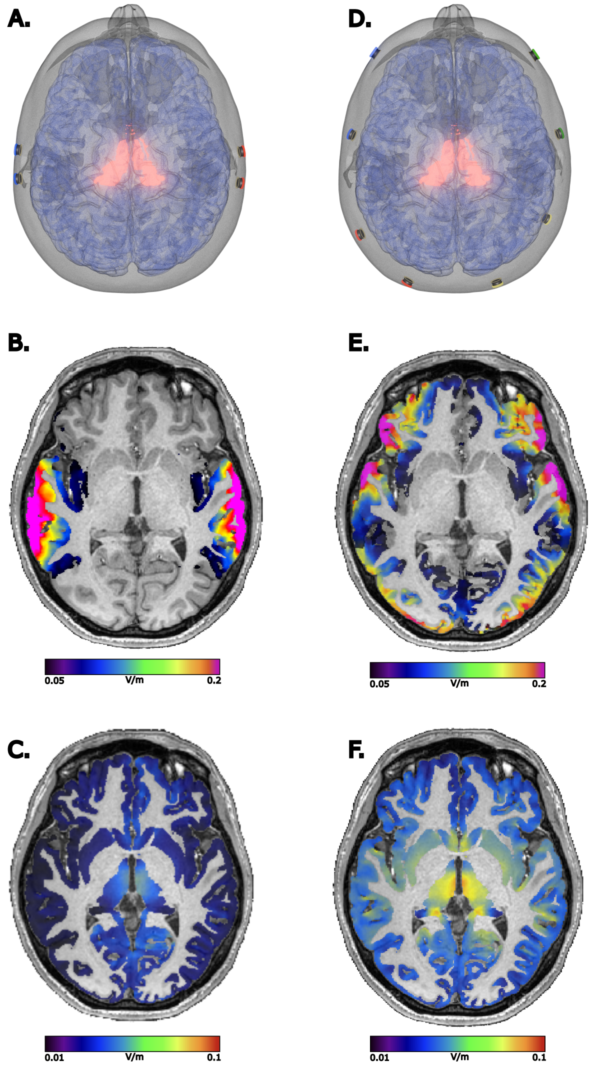


**Figure S2.** Multipolar vs Unipolar simulation Targeting the thalamus. **A.** ROI bi-lateral thalamus highlighted in red with two pairs of electrodes for traditional TI stimulation. **B.** Carrier fields on a scale of 0.05-0.2 V/m extract for the grey matter. **C.** TI field on a scale of 0.01 – 0.1 V/m. **D-F.** Similar panels for the mTI approach.

Focality optimization represents a critical challenge in TI stimulation, as it aims to maximize the ratio of field intensity within the target region relative to off-target areas, here defined as the rest of the grey matter. Unlike conventional optimization that maximizes absolute field strength, focality optimization explicitly penalizes field in the non-ROI, promoting spatial selectivity essential for precise neuromodulation. We evaluated three distinct thresholding approaches: fixed absolute thresholds (0.1 and 0.3 V/m) based on physiological relevance, and relative thresholds (50% and 80% of peak field) from a value previously recorded in the ROI with the goal of maximizing the mean $TI_{max}$.

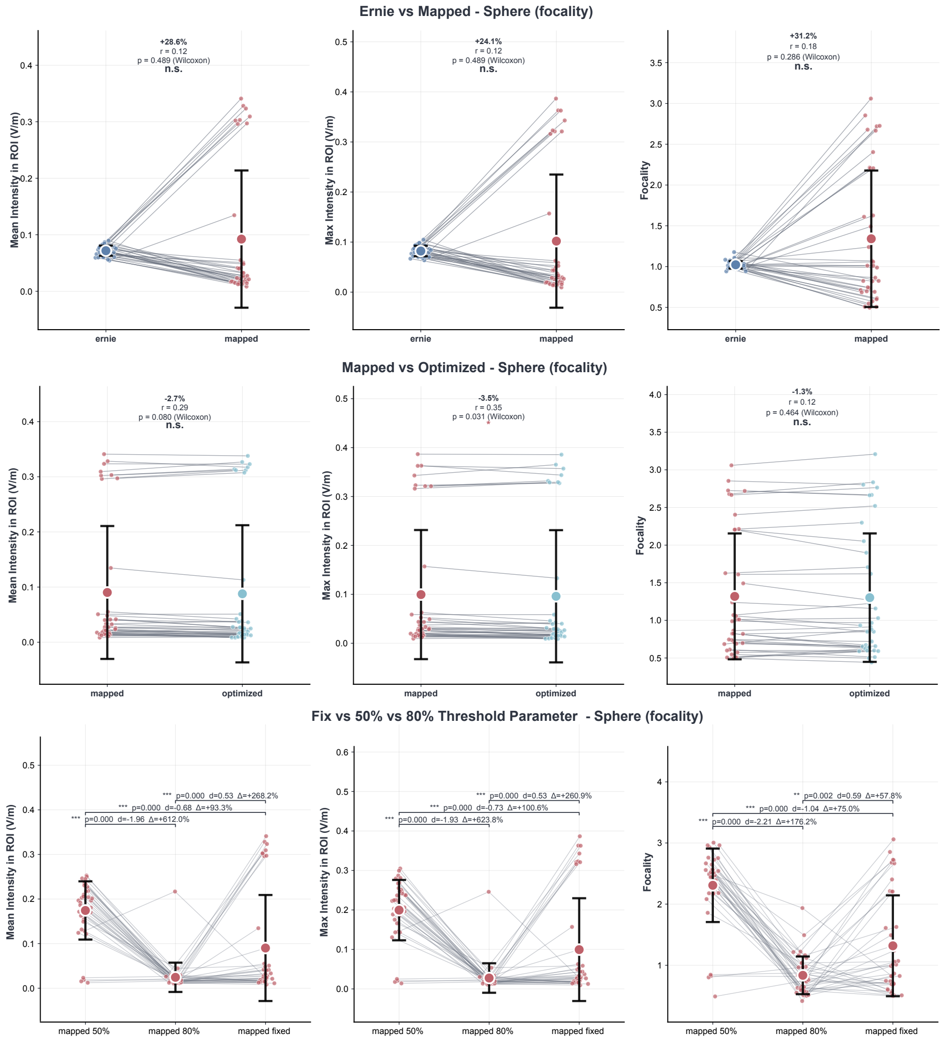


**Figure S3.** Comparative analysis of focality optimization under different thresholding strategies. **A.** Comparison of Ernie (generalized model) versus mapped electrode configurations using fixed thresholds (0.1 and 0.3 V/m). The mapped individualized montages achieved superior focality compared to the generalized approach (focality: 1.319 ± 0.814 vs 1.024 ± 0.045; ROI mean: 0.090 vs 0.072 V/m; gray-matter mean: 0.048 vs 0.070 V/m; n = 37), demonstrating that subject-specific electrode mapping enhances spatial selectivity by concentrating fields within the target while reducing off-target exposure though the results were not statistically significant. **B.** Evaluation of potential gains from unconstrained optimization beyond mapped solutions. The similar focality values between mapped and fully optimized montages (1.319 ± 0.814 vs 1.302 ± 0.831; n = 37) indicate that HD-EEG electrode constraints impose minimal penalty on achievable focality, supporting the practical viability of standardized electrode systems. **C.** Three-way comparison revealing the profound impact of threshold selection on focality metrics. Moving from fixed to 50% relative threshold increased focality by 75.0% ($p = 2.08\times10^{-6}$), while the 80% threshold showed 36.6% lower focality compared to the fixed threshold (all comparisons: Wilcoxon signed-rank test, $p < 0.01$).

The correlation matrix demonstrates the interrelationships between anatomical and demographic predictors of TI field variability. Strong within-tissue correlations between volume and thickness measurements necessitated collinearity handling in the multivariate analyses. Age showed weak to moderate correlations with tissue metrics, with the strongest associations observed with CSF volume (r = 0.294) and bone volume (r = -0.124). The high correlations between skin, bone, and CSF characteristics suggest that these anatomical features co-vary across individuals, potentially reflecting overall head size or morphological phenotypes. This correlation structure informed the variable selection strategy for the regression models, where volume metrics were retained over thickness metrics when both showed high correlation (|r| > 0.7) to avoid multicollinearity issues while maintaining predictive power.

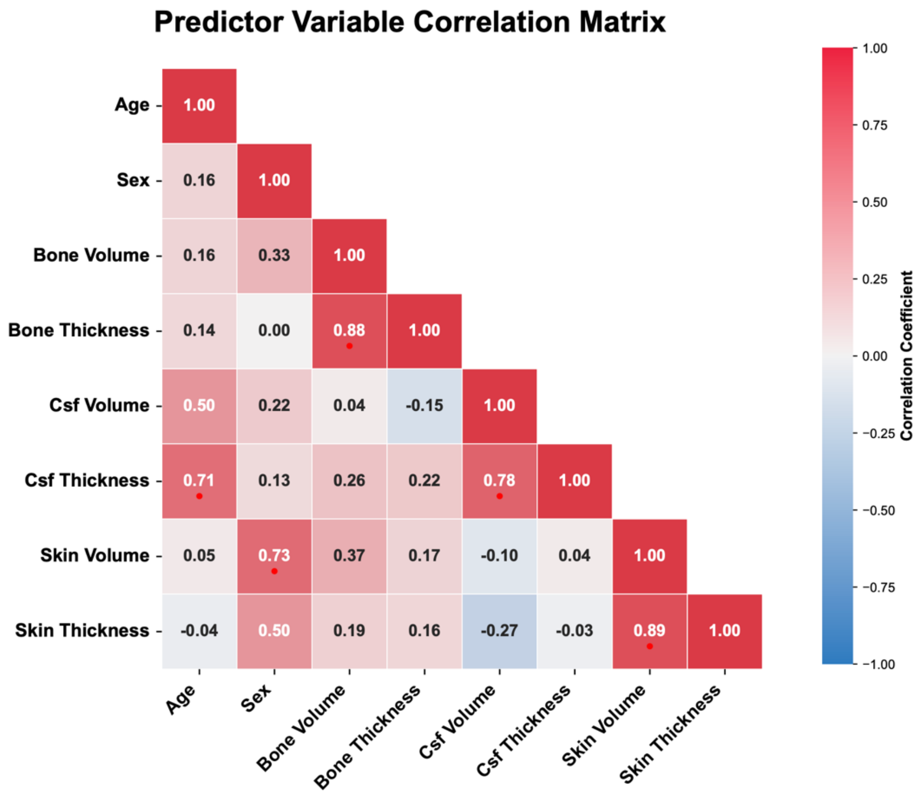


**Figure S4.** Correlation matrix of anatomical and demographic predictors. Correlation matrix showing Pearson correlation coefficients between anatomical metrics (bone volume, bone thickness, CSF volume, CSF thickness, skin volume, skin thickness) and demographic variables (age) across all subjects (n = 36). The matrix reveals strong positive correlations between volume and thickness measurements within each tissue type (bone: r = 0.883; CSF: r = 0.780; skin: r = 0.889), indicating multicollinearity that was addressed in the regression analyses by excluding thickness variables when correlations exceeded 0.7.

With the current implementation and the EEG net chosen in this study, the mapping function is less effective for the subcortical hippocampus target compared to the sphere/insula. This could be due to the larger separation required between electrodes and the limited coverage of the 185 HD-EEG net used in the lower density towards the back of the neck. The mapping distances for hippocampal targeting occasionally reached up to 30 mm, substantially larger than the 4-9 mm typical for cortical targets, suggesting that deeper structures may benefit from expanded electrode coverage or alternative montage strategies.

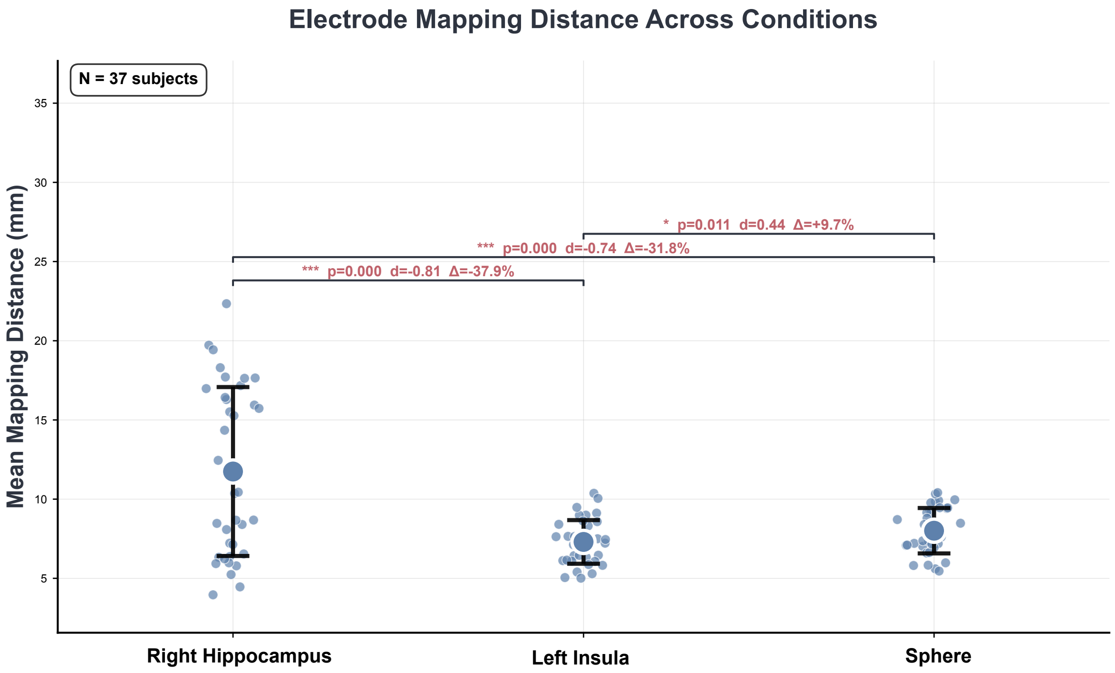


**Figure S5.** Electrode mapping distance comparison across anatomical targets. Within-subject analysis of mean Euclidean distances (mm) between optimized electrode positions and their mapped counterparts on the HD-EEG net for three distinct targets: right hippocampus (subcortical, 11.74 ± 5.33 mm), left insula (cortical, 7.30 ± 1.38 mm), and spherical ROI (36.10, 14.14, 0.33; radius 5 mm, 8.01 ± 1.43 mm). One-way repeated measures ANOVA revealed significant differences across targets ($F = 19.56$, $p < 0.0001$, $n = 37$). Post-hoc pairwise comparisons with Bonferroni correction showed the hippocampus required significantly larger mapping distances compared to both insula ($\Delta = 4.45$ mm, $p < 0.0001$) and spherical targets ($\Delta = 3.74$ mm, $p = 0.0002$), while the spherical target showed marginally larger distances than the insula ($\Delta = 0.71$ mm, $p = 0.033$). Individual participants shown as dots with slight horizontal jitter, connected by gray lines to illustrate within-subject patterns. Box plots display median, interquartile range, and outliers. The depth-dependent pattern of mapping distances reflects the challenge of targeting subcortical structures with scalp electrodes, where optimal montages require wider electrode separations that may exceed the local density of available HD-EEG positions.
